# An SH3-binding allosteric modulator stabilizes the global conformation of the AML-associated Src-family kinase, Hck

**DOI:** 10.1101/2024.11.05.622136

**Authors:** Ari M. Selzer, Gabriella Gerlach, Giancarlo Gonzalez-Areizaga, Thomas E. Wales, Stephanie Y. Cui, Prema Iyer, John R. Engen, Carlos Camacho, Rieko Ishima, Thomas E. Smithgall

## Abstract

While ATP-site inhibitors for protein-tyrosine kinases are often effective drugs, their clinical utility can be limited by off-target activity and acquired resistance mutations due to the conserved nature of the ATP-binding site. However, combining ATP-site and allosteric kinase inhibitors can overcome these shortcomings in a double-drugging framework. Here we explored the allosteric effects of two pyrimidine diamines, PDA1 and PDA2, on the conformational dynamics and activity of the Src-family tyrosine kinase Hck, a promising drug target for acute myeloid leukemia. Using ^1^H-^15^N HSQC NMR, we mapped the binding site for both analogs to the SH3 domain. Despite the shared binding site, PDA1 and PDA2 had opposing effects on near-full-length Hck dynamics by hydrogen-deuterium exchange mass spectrometry, with PDA1 stabilizing and PDA2 disrupting the overall kinase conformation. Kinase activity assays were consistent with these observations, with PDA2 enhancing kinase activity while PDA1 was without effect. Molecular dynamics simulations predicted selective bridging of the kinase domain N-lobe and SH3 domain by PDA1, a mechanism of allosteric stabilization supported by site-directed mutagenesis of N-lobe contact sites. Cellular thermal shift assays confirmed SH3 domain-dependent interaction of PDA1 with wild-type Hck in myeloid leukemia cells and with a kinase domain gatekeeper mutant (T338M). These results identify PDA1 as a starting point for Src-family kinase allosteric inhibitor development that may work in concert with ATP-site inhibitors to suppress the evolution of resistance.

## Introduction

Non-receptor protein-tyrosine kinases are diverse signaling molecules which affect a multitude of cellular processes in both normal and disease states. These enzymes share a well-conserved, bilobed kinase domain which transfers the γ-phosphate group from ATP onto select tyrosine residues of substrate proteins. The kinase active site is located at the interface of the two lobes and includes the flexible activation loop. While the kinase domain and ATP-binding site are highly conserved, the proteins have diverse regulatory domains N-or C-terminal to the kinase domain which can influence many kinase properties including dynamics, activity, localization and substrate specificity (1, 2).

Some of the best characterized examples of allosteric tyrosine kinase regulation come from the Src family of non-receptor tyrosine kinases (SFKs). There are eight mammalian SFKs: Src, Yes, Fyn, Hck, Lyn, Fgr, Lck, and Blk which can be separated into two four-member subfamilies (A and B) based on sequence similarity. Several family members, such as Yes and Src, are expressed in almost all cell types. Others exhibit lineage-specific expression patterns, such as Hck, which is expressed primarily in hematopoietic cells. In addition to the kinase domain, all SFKs have Src homology 2 (SH2) and SH3 domains, which are coupled by a short flexible connector. Other regulatory elements include the N-terminal SH4 myristoylation signal sequence, which is important for peripheral membrane localization, an unstructured unique region, the SH2-kinase domain linker, and a C-terminal tail (2, 3).

SFK activation is tightly controlled by intramolecular interactions as well as the phosphorylation state of the protein. In the inactive state, the SH3 domain binds to a polyproline type-II (PPII) helix formed by the SH2-kinase linker (Figure 1A) while the SH2 domain engages the C-terminal tail, which is phosphorylated by a separate regulatory kinase known as Csk (4). These interactions push the SH3-SH2 unit against the back of the kinase domain, allosterically stabilizing a closed, inactive conformation (Figure 1A). Interactions with ligands that perturb these interactions as well as dephosphorylation of the regulatory tail phosphotyrosine, pTyr527 (amino acid numbering based on the crystal structure of human Src; PDB: 2SRC) (5), cause the protein to become activated. Full kinase activation also involves autophosphorylation of the activation loop tyrosine (pTyr416). This phosphorylation event creates a steric and electrostatic barrier that prevents the activation loop from adopting a partially folded, inactive conformation. Additional amino acids, such as Trp260, affect these regulatory events which are not mutually exclusive and the formation of one activating or inactivating interaction can propagate to others (6). SFKs can also adopt alternative active forms in which either the SH3 or SH2 domains are displaced from their regulatory interactions (7, 8). In addition to regulating kinase activity, the non-catalytic regions modulate signal transduction by interacting with SH3 and SH2 binding motifs in other proteins to regulate substrate selection and subcellular localization. Less is known about the functions of the unique domains, which vary in length and sequence between family members. NMR studies suggest that the unique region may interact with the SH3 domain to regulate kinase activity as well as substrate selection (9).

**Figure 1.**
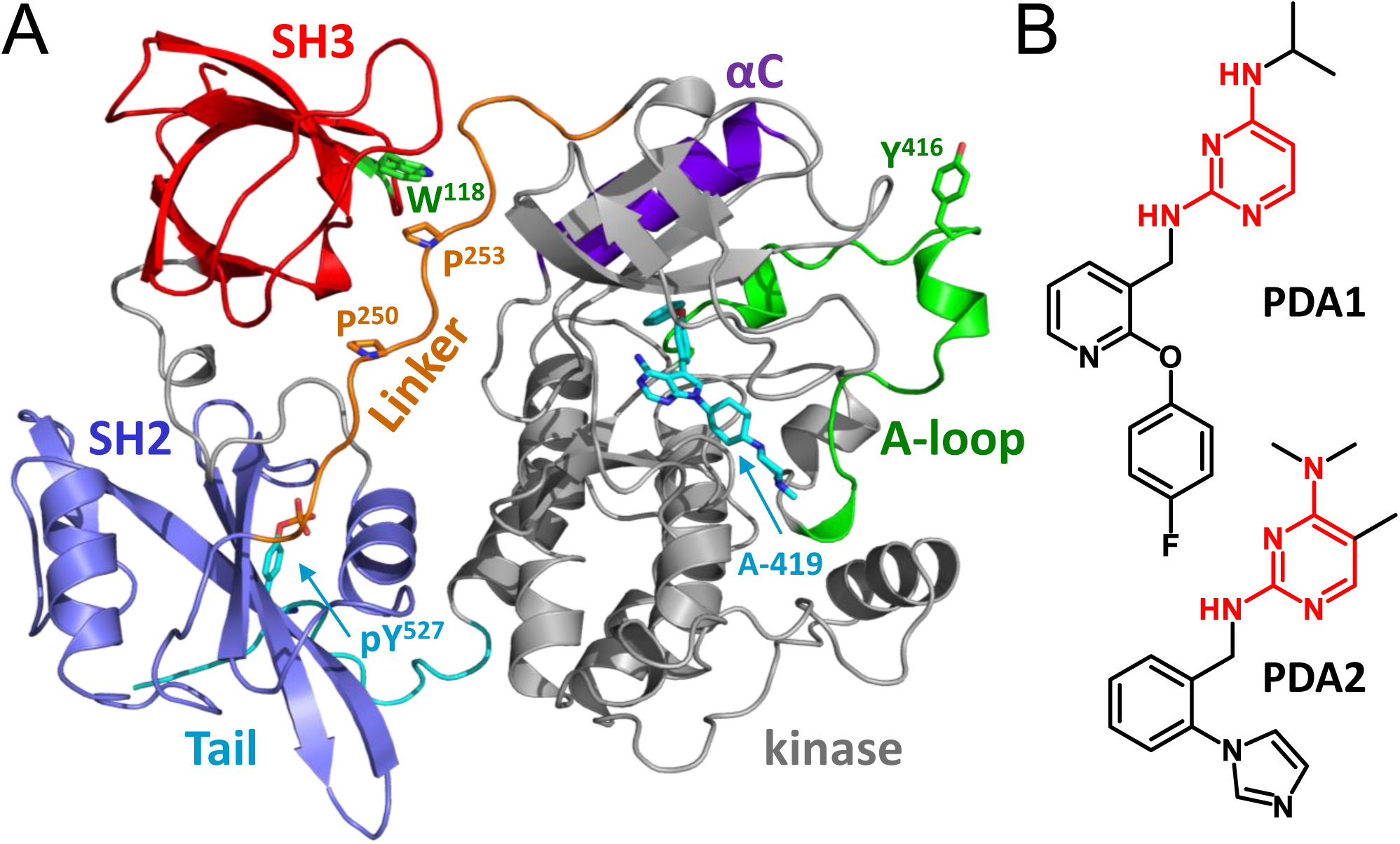
Crystal structure of near-full-length Hck and chemical structures of pyrimidine diamine allosteric modulators. A) Crystal structure of Hck bound to the ATP-site inhibitor, A-419259 (A-419; cyan). In the presence of this inhibitor, Hck adopts the closed conformation with the SH3 domain (red) bound to the SH2-kinase linker (orange) and the SH2 domain (blue) engaging the phosphorylated C-terminal tail (cyan; pTyr527). Highlighted features of the kinase domain include the C-helix (purple) in the N-lobe as well as the activation loop (green) and autophosphorylation site (Tyr416). The SH2-kinase linker forms a polyproline type II helix involving prolines 250 and 253 (side chains shown) which engages the SH3 domain; SH3 Trp118 (green) is essential for this interface to form. Model produced using PyMol (Schrödinger) and Hck crystal coordinates from PDB entry 9BYJ. B) Chemical structures of PDA1 and PDA2. The common pyrimidine diamine scaffold is highlighted in red.

Tyrosine kinases can become oncogenic when not properly regulated. The fusion protein Bcr-Abl, which results from the Philadelphia chromosome translocation, is the oncogenic tyrosine kinase driver of chronic myeloid leukemia (CML). The Abl portion of Bcr-Abl is a tyrosine kinase that is structurally related to the SFKs, with an SH3-SH2-kinase core followed by a long C-terminal region (10). The N-terminal 70 amino acids of Bcr form a coiled-coil domain which induces oligomerization and constitutive activation of Abl in this fusion protein (11, 12). Imatinib, a selective Abl kinase inhibitor, received FDA approval for use in CML in 2001 and inspired a new generation of kinase-directed drug discovery (13). At the beginning of 2024, 80 small molecule kinase inhibitors had received FDA approval, 63 of which inhibit tyrosine kinases including SFKs (14).

Because SFKs normally control cellular growth, differentiation, adhesion and motility, dysregulation of SFK activity contributes to many forms of cancer (15, 16). Of note, the myeloid cell specific SFKs, Fgr, Hck, and Lyn, are associated with both chronic and acute (AML) myeloid leukemias. AML is the most common form of blood cancer in adults with approximately 20,000 new cases and 11,000 deaths in the United States each year (17). High-level expression of Fgr, Hck, or Lyn correlates with poor prognoses for AML patients (18). Hck specifically is overexpressed in leukemia stem cells compared to healthy hematopoietic stem cells, making it a particularly attractive target for inhibitor discovery (19, 20). Several preclinical ATP-site inhibitors with nM activity against Hck have been described, including the pyrrolopyrimidine A-419259 (20) and the N-phenylbenzamide TL02-59 (21). Both compounds block the growth of AML cells that overexpress these kinases in experiments with cell lines, patient bone marrow cells, and mouse models of the disease (20, 21).

Despite the promise of ATP-site tyrosine kinase inhibitors (TKIs) in cancer, off-target activity and acquired resistance remain important clinical limitations. Most clinical TKIs target the ATP-binding-site which is highly conserved across different protein tyrosine kinase families. While some degree of selectivity can be achieved by targeting unique sites adjacent to the ATP-binding pocket, off-target binding can still lead to significant side effects. Additionally, a single point mutation in the drug binding site is often sufficient to cause loss of inhibitor activity. A common example involves the so-called ‘gatekeeper’ residue (Thr315 in Abl, Thr338 in c-Src), which when mutated can cause resistance for many TKIs including imatinib (22) while also increasing kinase activity (23). As a result, recent TKI discovery has focused on allosteric inhibitors which target sites adjacent to the ATP-binding site or more distant pockets and have the potential to work in combination with ATP-site inhibitors (24).

To overcome the issue of acquired resistance, a “double-drugging” approach has emerged which involves the simultaneous use of two inhibitors for the same kinase which have mutually exclusive resistance mechanisms. Wylie *et al*. (25) demonstrated this concept for CML by combining the FDA-approved Abl inhibitors nilotinib, which binds the ATP-site, and asciminib, which targets a unique myristoyl binding pocket in the C-lobe of the Abl kinase domain. When used separately, both inhibitors initially suppressed CML cell growth in a mouse model, followed by relapse presumably due to acquired resistance mutations within the drug binding site. However, administration of the two-drug combination resulted in a durable response, with no tumor regrowth even after dosing was stopped. In a related study, Kim *et al*. (26) showed that these two inhibitors increase each other’s affinity when they have the same conformational preferences. This “double drugging framework” occurs because binding of one inhibitor shifts the conformational landscape of the kinase domain to favor the preferred conformation of the second inhibitor, thereby decreasing the thermodynamic barrier for binding. In practice, this approach has the potential of increasing the specificity of the inhibitor combination by increasing the difference between their on-vs. off-target affinities.

In the present study, we evaluated the binding mechanism and conformational influence of two small molecules that interact with the regulatory SH3-SH2-linker region of the myeloid SFK, Hck. These compounds, PDA1 and PDA2, share a pyrimidine diamine scaffold (Figure 1B) and are shown here to bind to the PPII helix-interacting site of the Hck SH3 domain by ^1^H-^15^N HSQC NMR. Despite the shared binding site, these compounds have opposing effects on the overall kinase conformation as shown by hydrogen-deuterium exchange mass spectrometry (HDX-MS). Molecular dynamics (MD) simulations and site-directed mutagenesis suggest that PDA1, which stabilizes kinase dynamics, bridges the SH3 domain and the N-lobe of the kinase domain, allowing the compound to stabilize a closed, downregulated conformation. PDA2, on the other hand, destabilizes the overall kinase conformation and stimulates kinase activity in vitro. Using a cellular thermal shift assay (CETSA), which provides information on overall protein stability in cells, we demonstrate that PDA1 interacts with Hck in myeloid leukemia cells including a gatekeeper mutant which confers resistance to the ATP site inhibitor, A-419259. Conversely, an SH3 mutant of Hck that does not bind PDA1 (W118A) retains binding to the orthosteric inhibitor, A-419259. Development of PDA1 analogs with improved potency for interdomain bridging and cellular activity represents a promising path forward for allosteric inhibitors of Hck and other SFKs. These inhibitors may have therapeutic efficacy by themselves or as part of a combination therapy with ATP-site inhibitors in a double-drugging framework for AML cases where Hck is overexpressed.

## Results

### Determination of PDA binding site on the SH3 domain of Hck by HSQC NMR

We previously reported the identification of small molecules that bind to the non-catalytic SH3-SH2-linker region of Hck (27). This effort identified a cluster of small molecules that share a common pyrimidine diamine (PDA) core from a fluorescence polarization-based screen of 60,000 diverse compounds. Each of these compounds bound to the Hck regulatory region with micromolar potency by surface plasmon resonance (SPR). To begin to define the binding site and mechanism of action, we first resynthesized two of the compounds (PDA1 and PDA2) and confirmed their structures by NMR and LC-MS (synthetic routes and analytical data are provided in the Supplemental Information).

To experimentally determine the binding site of the PDA compounds within the Hck regulatory region, an ^15^Nlabeled Hck SH3-SH2-linker (Hck-32L) protein was prepared for analysis by ^1^H-^15^N HSQC NMR. Previous NMR studies of a similar Hck-32L protein identified backbone amide N-H resonances (28), which enabled identification of spectral changes in response to PDA binding. Both PDA1 and PDA2 induced concentration-dependent chemical shift perturbations (CSPs) in resonances which correlate to the backbone amides of SH3 residues including Glu94, Trp118, Trp119, Leu124, and Ile132. Other CSPs were associated with SH3-SH2 connector residues Thr145 and Phe150 as well as Glu255 in the SH2-kinase linker (Figure 2A and 2B). Complete Hck-32L spectra in the presence of each compound are provided in the Supplemental Information (Figure S1).

**Figure 2.**
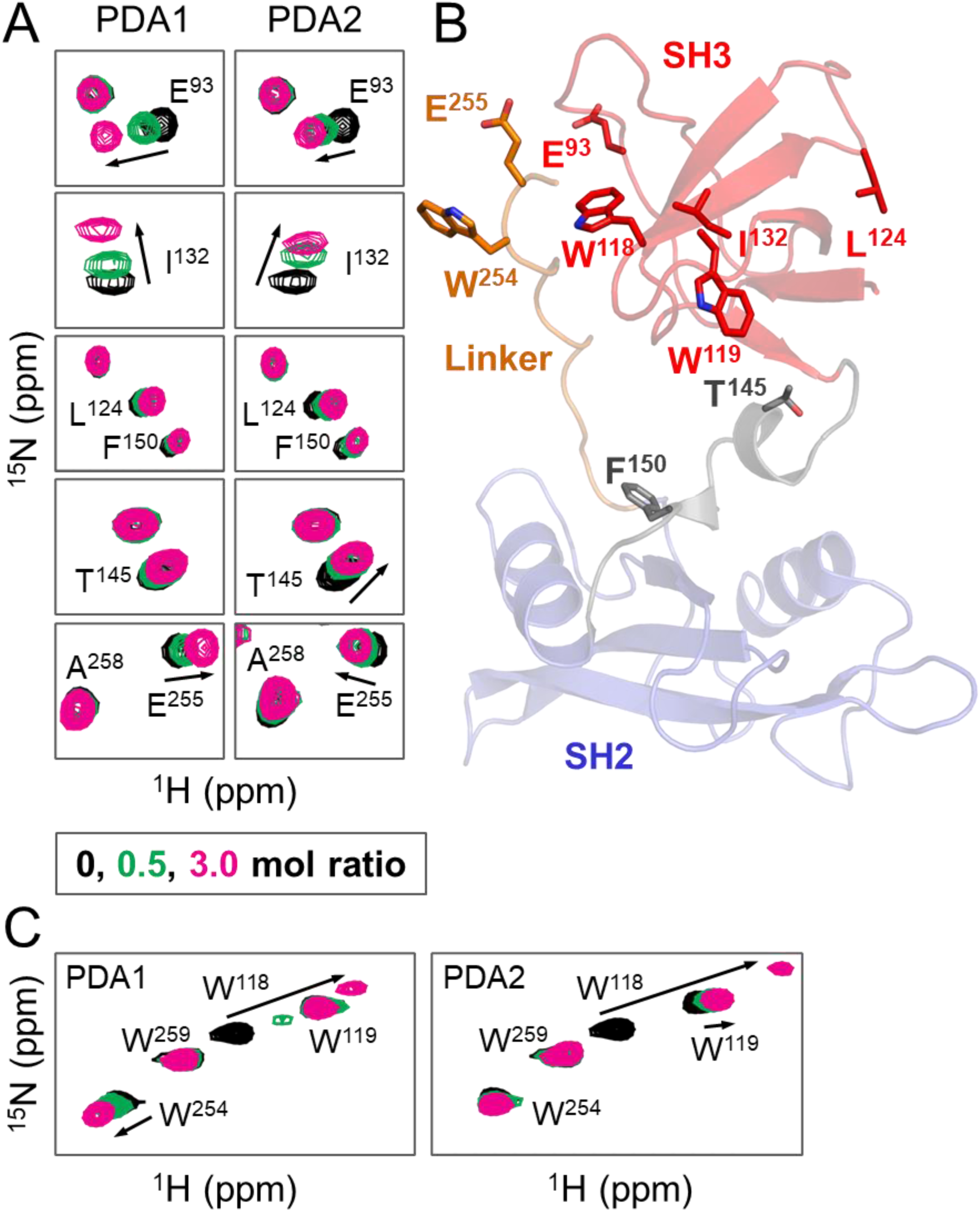
HSQC NMR analysis of Hck SH3-SH2-linker protein interaction with PDA analogs. PDA1 and PDA2 were titrated with ^15^N-labeled Hck-SH3-SH2-linker protein and ^1^H-^15^N HSQC NMR spectra were recorded. Backbone amide resonances were identified from a previous assignment (28). Tryptophan indole amide resonances were assigned by mutagenesis (Figures S2, S3 and S4). Significant chemical shift perturbations (CSPs) observed in a subset of (A) backbone amides and (C) tryptophan indole amides are mapped on the crystal structure of the Hck SH3-SH2-linker region in panel B. CSPs induced by PDA binding observed in the SH3 domain include E93, W118, I132. PDA1 shows more pronounced CSPs of amides, compared to PDA2, in the connector between SH3 and SH2 (E144 and T145) and W254 and E255 in the SH2-kinase linker. SH3-SH2-linker protein concentration in this experiment was 60 µM.

Additional insight regarding the binding site is provided from analysis of CSPs involving tryptophan indole N-H resonances in the SH3 domain (Trp118 and 119) as well as the SH2-kinase linker (Trp254; Figure 2C). Both PDA1 and PDA2 induced large CSPs of the indole N-H resonance of Trp118. Previous studies have shown that Trp118 is essential for Hck SH3 binding activity both in vitro and in cells (27, 29). Neither compound strongly affected the indole N-H resonance of adjacent Trp119. In crystal structures of Hck, the Trp118 side chain forms part of a surface that interacts with the PPII helix in the linker, while the Trp119 side chain faces the hydrophobic core of the domain (30). This suggests that the PDA compounds may bind near Trp118 in the PPII helix interaction site. To unambiguously assign the Trp118 indole N-H resonance, HSQC NMR experiments were repeated with an Hck-32L-W118A mutant protein. Spectra with this mutant showed loss of the Trp118 signal as well as other CSPs associated with binding (Figure S2). Finally, PDA1 but not PDA2 induced a CSP in Trp254 in the linker, providing evidence that each ligand has a unique effect on the overall conformation of the regulatory region. Because there are two tryptophan residues in the linker (Trp254 and Trp259), the PDA-induced Trp254 indole resonance was unambiguously assigned with an Hck-32L-W254A mutant (Figure S3A). Control spectra show minimal effects of the DMSO co-solvent on the HSQC spectra of the Hck SH3-SH2 protein (Figure S3B).

The initial report describing the discovery of the PDA compounds showed that the SH3 domain alone was not sufficient for binding by SPR (27). However, the NMR results presented above show that the SH3 domain alone likely provides the key site for PDA binding. To determine whether NMR can detect PDA binding to the SH3 domain alone, ^15^N-labeled Hck SH3 protein was produced, and ^1^H-^15^N HSQC spectra were recorded in the presence and absence of each PDA compound. CSPs observed with the isolated SH3 domain in the presence of each compound were nearly identical to those observed with the Hck-32L protein, indicating that the SH3 domain is the primarily interaction site of PDAs (Figure S4). Detailed NMR titration data with each PDA compound revealed that the major CSPs map to the binding surface of the SH3 domain (Figure S5A and S5B). Titration curves for the CSPs associated with three residues in this region (Glu94, Ala121, Trp118) show saturation and yielded dissociation constants of 42.6 ± 4.7 µM for PDA1 and 65.1 ± 3.1 µM for PDA2 (Figure S5C and S5D).

### PDA compounds show reduced binding affinity for other Src-family kinases by SPR

Establishing on-target selectivity, especially within a conserved kinase family, is a challenge for ATP-site kinase inhibitor development. By targeting the SH3 domain, however, PDA compounds may have the potential for improved selectivity compared to ATP-site inhibitors for Src-family members. To address this issue, we first aligned the sequences of the eight human SFKs and calculated the amino acid sequence identity compared to Hck. The average overall sequence identity of Hck to all other family members was 63.2% with a standard deviation of 4.8% (Figure 3A). We then calculated the identity for the individual SH3, SH2, and kinase domains as well as the ATP-binding sites. Not surprisingly, the kinase domains and ATP binding sites showed the highest conservation with Hck (72.6 ± 5.9% and 83.9 ± 5.9% identity, respectively). Interestingly, every single kinase domain and ATP-binding site showed higher conservation to those in Hck compared to the full-length protein, supporting evolutionary pressure against mutations in this region. The opposite trend was observed for the SH3 domains, which were the least conserved domains overall compared to Hck (56.0 ± 4.5% identity with Hck SH3), and each SFK has higher overall sequence identity to Hck than the SH3 domains have to one another. For the SH2 domains, two were less conserved, three were more conserved, and two were about the same, with 56.0 ± 4.5% average identity with Hck.

**Figure 3.**
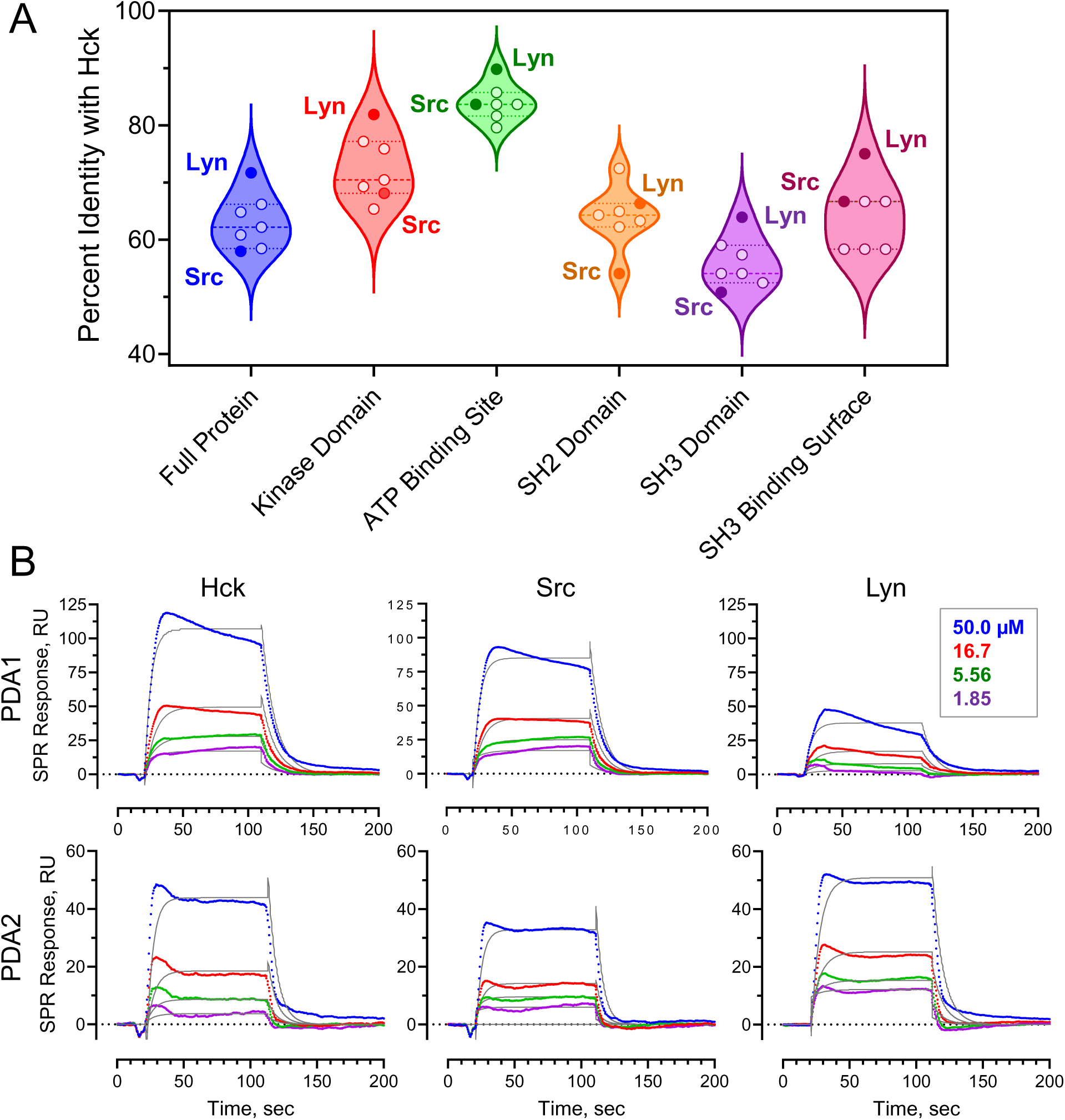
Sequence identity among SFK domains and binding selectivity of PDAs to Hck, Src and Lyn. A) Violin plot showing distribution of sequence identities among Src family members and their domains relative to Hck. Lyn is the closest relative to Hck, while Src is among the most distant with the SH3 domains showing the least sequence conservation. B) SPR analysis of PDA1 and PDA2 binding to near-full-length Hck, Src and Lyn. Each kinase protein was immobilized on the SPR chip, and the PDA analogs were injected over the range of concentrations shown in triplicate. Representative sensorgrams are shown in color with fitted curves overlaid in black. K_D_ values are reported in the text.

We also explored conservation in the binding site of the SH3 domain across the SFKs (Figure 3A). To define this region, we chose SH3 residues within 5 Å of the polyproline helix of the linker in a closed structure of Hck (PDB: 9BYJ) (31). These include Y90, D91, I95, H96, E117, W118, Y131, P133, N135, and Y136. These residues were then compared to residues in the analogous positions in the SH3 domains of the other SFKs. The Hck SH3 binding surface shows a similar degree of conservation as the overall kinases (63.2% vs 64.3%) but lower conservation compared to the kinase domains (72.6%) and the ATP-binding site (83.96%). Thus, enough sequence diversity may be present to create PDA analogs with isoform selectivity.

We next explored the binding of PDA compounds to other members of the Src kinase family to investigate whether sequence differences in the SH3 domains might influence PDA binding selectivity. For these experiments we chose Lyn, in which the SH3 domain is most closely related to Hck, and Src, which is more distantly related (Figure 3A). These kinases also represent members of the two SFK subfamilies, type A (Src) and type B (Lyn and Hck). Interaction kinetics were assessed by SPR using recombinant near-full-length Hck, Lyn and Src. All three recombinant proteins lack the N-terminal unique domain, are phosphorylated on their C-terminal tail by co-expression of Csk and dephosphorylated on the A-loop by co-expression of the PTP1B catalytic domain. As a result, each kinase adopts the closed conformation. In all cases, the PDA compounds showed the highest affinity for Hck. For PDA1, the K_D_ value for Src (55.0 ± 2.18 µM) was slightly higher than for Hck (42.2 ± 2.88 µM), while with Lyn, PDA1 showed a K_D_ value (74.9 ± 19.9 µM) nearly twice that of Hck (Figure 3B). The replacement of Hck Glu93 and Glu255, both of which showed significant CSPs by NMR, with shorter aspartate side chains in Lyn may impact the binding of PDA1 despite charge conservation. With PDA2, both Lyn and Src showed higher K_D_ values (180 ± 33.7 µM and 236 ± 18.1 µM, respectively) than Hck (110 ± 24.7 µM). While the differences in the K_D_ values among the kinases tested is modest, it is important emphasize that these differences were observed with hit compounds from a screen that have not undergone optimization. Analysis of sequence conservation in the SH3 binding surface described above suggests that structure-based design may ultimately yield analogs with improved selectivity and specificity. That said, some cross-over in selectivity within the Src family may be of benefit, because multiple Src family members have been implicated in etiology of AML. In addition to Hck, these include Fgr, Fyn and Lyn (18).

### PDA1 and PDA2 have opposite effects on Hck conformational dynamics and kinase activity

Previous studies have shown that peptide and protein ligands for the Hck SH3 domain stimulate kinase activity both *in vitro* and in cells (32-35). These observations support a mechanism by which SH3 domain displacement leads to allosteric activation of the kinase domain. Because the PDA compounds bind to the SH3 domain, we explored their impact on Hck conformational dynamics by hydrogen-deuterium exchange mass spectrometry (HDX-MS). The first series of experiments involved the Hck-U32L protein, which lacks the kinase domain and C-terminal tail but retains the N-terminal unique domain. Hck-U32L was preincubated in the presence or absence of each PDA analog followed by exposure to D_2_O-based buffer to induce deuterium exchange for various times ranging from 10 seconds to 4 hours. Following exchange, deuterium uptake into peptic peptides derived from the protein was determined by LC-MS/MS. Overall results are presented as difference heat maps in terms of protection from (reduced HDX, blue) or exposure to (increased HDX, green) deuterium uptake as a function of D_2_O exposure time in the presence or absence of the compound (Figure 4). Incubation with either PDA1 or PDA2 caused an overall decrease in HDX for peptides derived from the SH3 domain, including peptide with residues exhibiting significant CSPs in the NMR studies. These include peptide 109-135, which contains Trp118 that was identified as a key interaction point by mutagenesis (individual deuterium uptake curves for this peptide are shown in Figure 5A). This observation supports the conclusion that PDA1 and PDA2 occupy overlapping binding sites in the SH3 domain and are consistent with the NMR results. PDA1 also induced rapid and sustained protection of the unique region, as well as the SH2 domain and the SH2-kinase linker (Figures 5B-D). By contrast, PDA2 had no allosteric influence on deuterium uptake by the unique, SH2, and linker regions, despite the shared binding site with PDA1. These results suggest that PDA1 acts as an allosteric stabilizer of the Hck regulatory region, an unexpected result given the binding site of the compound.

**Figure 4.**
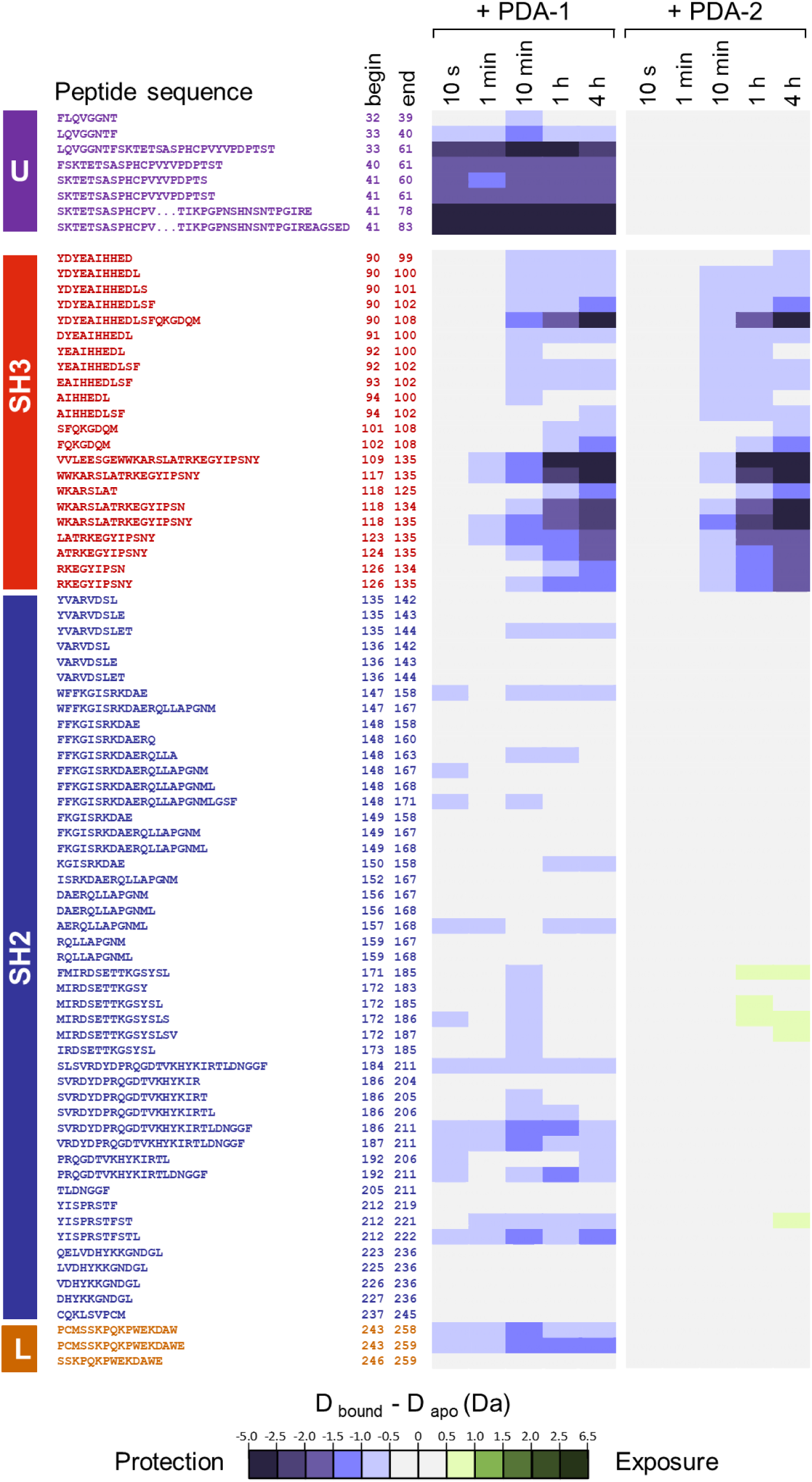
HDX-MS difference maps for Hck-U32L protein in the presence of PDA1 and PDA2. Hck-U32L protein was equilibrated in the presence and absence of PDA1 and PDA2, followed by exposure to D2O based buffer for the timepoints shown. Peptic peptides were identified by LC-MS/MS and changes in deuterium uptake are shown as colored squares according to the scale. Peptide locations within the protein are indicated at left. All values used to make these heat maps are provided in the HDX Supplementary Data File.

**Figure 5.**
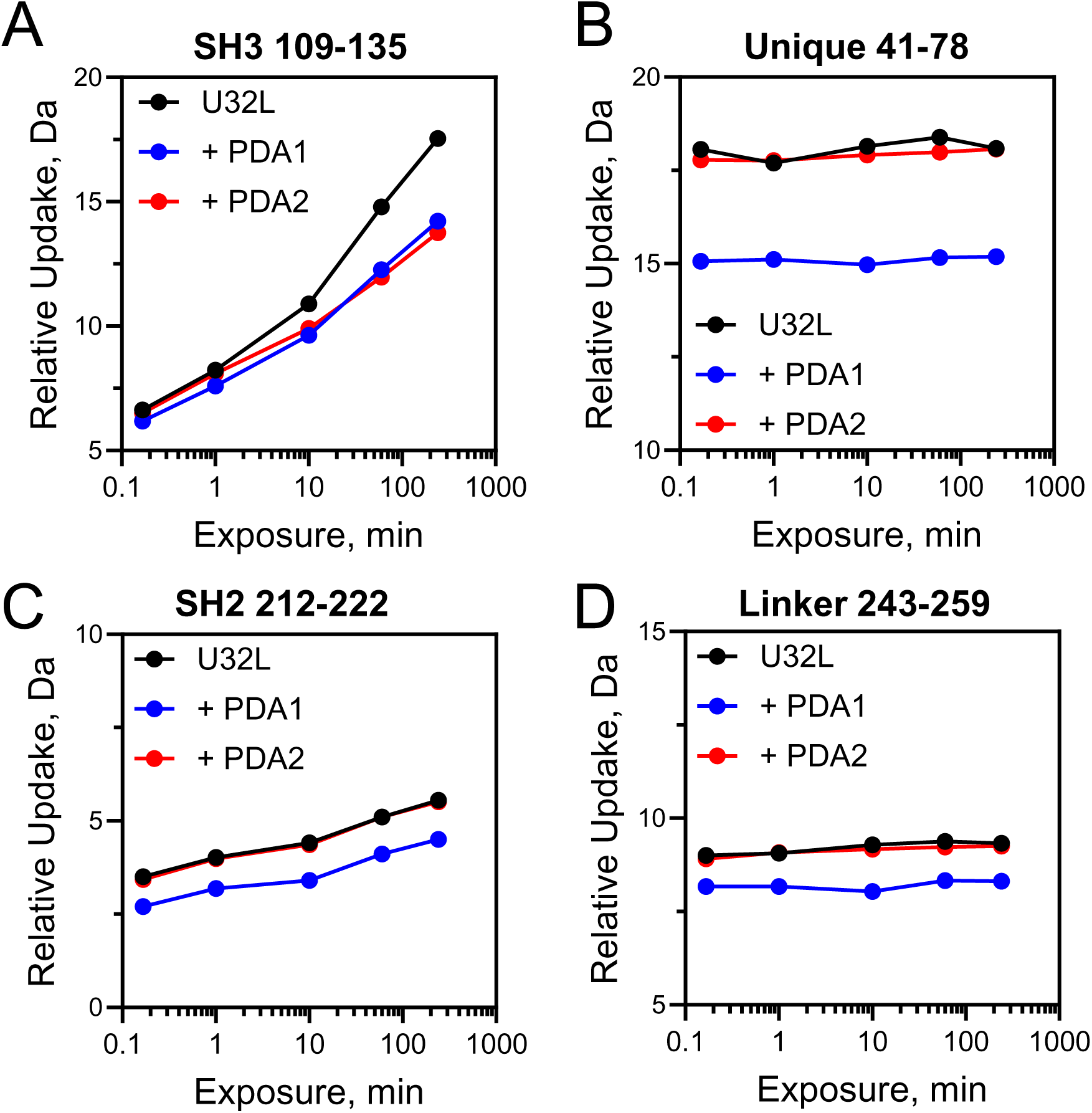
Deuterium uptake plots for Hck-U32L peptic peptides. The Hck-U32L protein was equilibrated in the absence or presence of PDA1 or PDA2 followed by HDX-MS analysis as shown in Figure 4. Deuterium uptake curves are shown for peptides derived from the SH3 domain (A), the N-terminal unique domain (B), the SH2 domain (C), and the SH2-kinase linker (D). All values used to make these heat maps are provided in the HDX Supplementary Data File.

We next performed HDX-MS analysis of recombinant near-full-length Hck, which lacks the unique domain, retains the kinase domain and C-terminal tail, and adopts the closed conformation. To facilitate visual comparison of regional differences in deuterium uptake in the presence of PDA1 and PDA2, combined HDX changes are mapped to a crystal structure of Hck (Figure 6A). Complete difference heat maps for all peptic peptides that were followed in the deuterium exchange experiment with near-full-length Hck are shown in the Supporting Information (Figures S6A and S6B and the HDX Supplemental Data File). Overall, PDA1 increased protection from deuterium exchange across diverse regions of the protein, including the SH3 and SH2 domains, the SH3-SH2 domain connector, and the kinase domain C-lobe. One exception is the SH2-kinase linker. While several peptides from this region showed an increase in deuterium uptake at earlier time points (10 s, 1 and 10 min) relative to the apo protein, they ultimately did not reach the same level of incorporation by the 4-hour time point. Deuterium uptake plots for linker peptide 243-267 illustrate this effect (Figure 6B). This observation is consistent with small molecule binding at the SH3 domain affecting the linker conformation or by directly inhibiting deuterium exchange through steric effects. In addition, a few peptides derived from the kinase domain N-lobe showed a modest increase in deuterium uptake at the longest time points tested (1 and 4 h); uptake curves with peptide 281-300 provide an example (Figure 6C).

**Figure 6.**
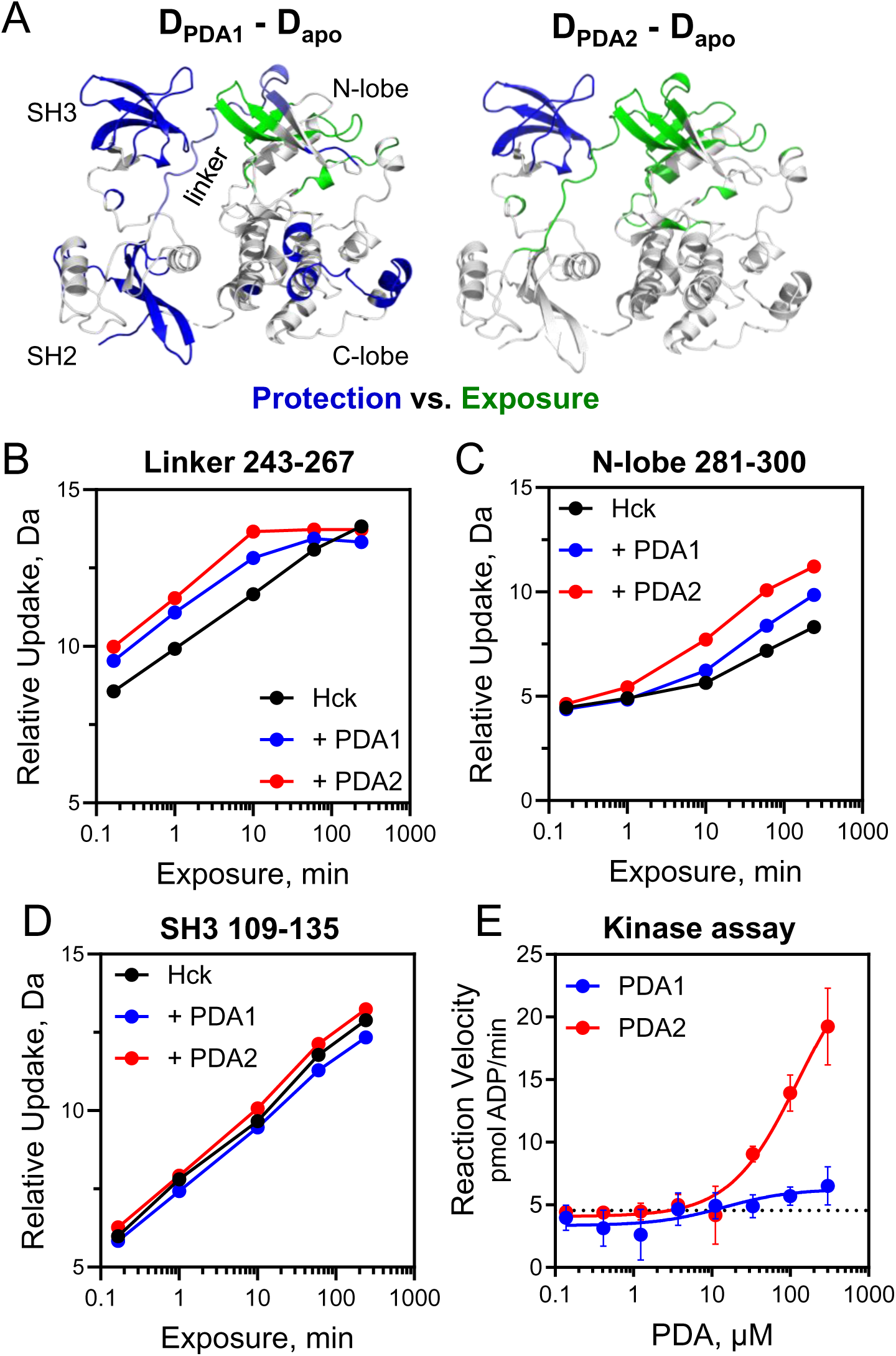
PDA1 and PDA2 have opposing effects on Hck dynamics and kinase activity. Recombinant near-full-length Hck was equilibrated with PDA1 or PDA2 or left untreated followed by HDX-MS analysis. Complete HDX-MS peptide deuterium uptake difference maps are presented in Figure S6 and in the HDX Supplementary Data File. A) Peptides showing changes in protection from (blue) or exposure to (green) deuterium uptake in response to PDA treatment are mapped on a crystal structure of Hck (PDB: 9BYJ). HDX data from the U32L protein and the kinase domain from nearfull-length Hck were combined to show the overall impacts of each PDA analog. Any peptide showing a significant difference in deuterium uptake in the presence of ligand at the 4 h time point is highlighted as shown. Individual deuterium uptake curves are shown for peptides derived from the SH2-kinase linker (B), the N-lobe of the kinase domain (C), and the SH3 domain (D). Kinetic kinase assays were performed with the same preparation of Hck used for the HDX-MS analysis (E). Reaction velocities were measured over the range of PDA1 and PDA2 concentrations shown.

Unlike PDA1, the addition of PDA2 to near-full-length Hck resulted in increased deuterium incorporation in peptides spanning the SH3-SH2 connector, and localized regions in both lobes of the kinase domain (Figure 6A). PDA2 also induced changes in the SH2-kinase linker consistent with displacement from the SH3 domain and to a greater degree than PDA1 (peptide 243-267; Figure 6B). The increased level of deuterium incorporation induced by PDA2 for the N-lobe peptide 281-300 reflects an increase in overall dynamics for this region when compared to both the PDA1-bound and apo proteins (Figure 6C). Overall, these changes in HDX suggest that PDA2 destabilizes and shifts Hck away from the closed conformation.

Because the Hck SH3 and SH2 domains allosterically regulate kinase activity, the results of the HDX-MS experiments predict that PDA1 may be an inhibitor of kinase activity while PDA2 may be an activator. To test this idea, we performed kinetic kinase activity assays with near-full-length Hck, the same recombinant kinase protein used in the HDX-MS studies. Kinase reactions were run over a wide range of PDA concentrations, and steady state reaction velocities were calculated by regression analysis of the linear part of each progress curve. Consistent with the HDX-MS results, PDA2 stimulated Hck activity in a concentration-dependent manner (Figure 6E). PDA1, however, had no significant effect on substrate peptide phosphorylation by Hck in this assay over the same range of concentrations (Figure 6E). In some experimental replicates, however, we observed concentration-dependent inhibition of Hck activity by PDA1 with micromolar IC_50_ values in line with those observed for PDA1 binding by SPR (data not shown). The low affinity of PDA1 for Hck as well as possible heterogeneity in the starting conformation of the kinase may account for the inconsistency in these results.

### Molecular dynamics simulations predict distinct interactions for PDA1 vs. PDA2

To investigate how PDA1 and PDA2 share a binding site yet have opposite effects on Hck dynamics and kinase activity, we performed MD simulations. The compounds were docked to a crystal structure of near-full-length Hck using the program Smina (36) to determine a starting conformation of the compounds before running 600 ns MD simulations. Movies of the complete simulations are provided in the Supporting Information. While PDA1 quickly settled into a preferred binding conformation, PDA2 was highly mobile at the beginning of the simulation, stably binding only after conformational changes to the SH3 domain as described below.

Models from the final frame of each simulation are shown in Figure 7A. Both PDA1 and PDA2 are positioned near SH3 domain Trp118 that is a key binding determinant identified by NMR in terms of CSP (Figure 2, S2, S4 and S5). However, their respective pyrimidine diamine scaffolds are orthogonally inverted relative to one another with different binding modes. PDA1 forms a series of interactions that bridge the SH3 domain and the N-lobe of the kinase domain, involving N-lobe residues His289 and Thr290. The difluorophenyl ring of PDA1 inserts between the side chains of His96 in the SH3 domain RT loop and Trp118 to form an ‘aromatic sandwich’. Additional contacts are present with SH3 residues Tyr92 and Tyr136. Binding of PDA2, on the other hand, required restructuring of the SH3 domain RT-loop. The positions of both small molecules adjacent to the binding surface of the SH3 domain suggest that they must displace the SH2-kinase linker, which has been previously associated with kinase activation as described above. However, only PDA2 induces kinase activation, suggesting that PDA1-mediated bridging of the SH3 domain and the N-lobe may compensate for linker displacement, resulting in the observed stabilization of the overall structure by HDX-MS and lack of kinase activation.

**Figure 7.**
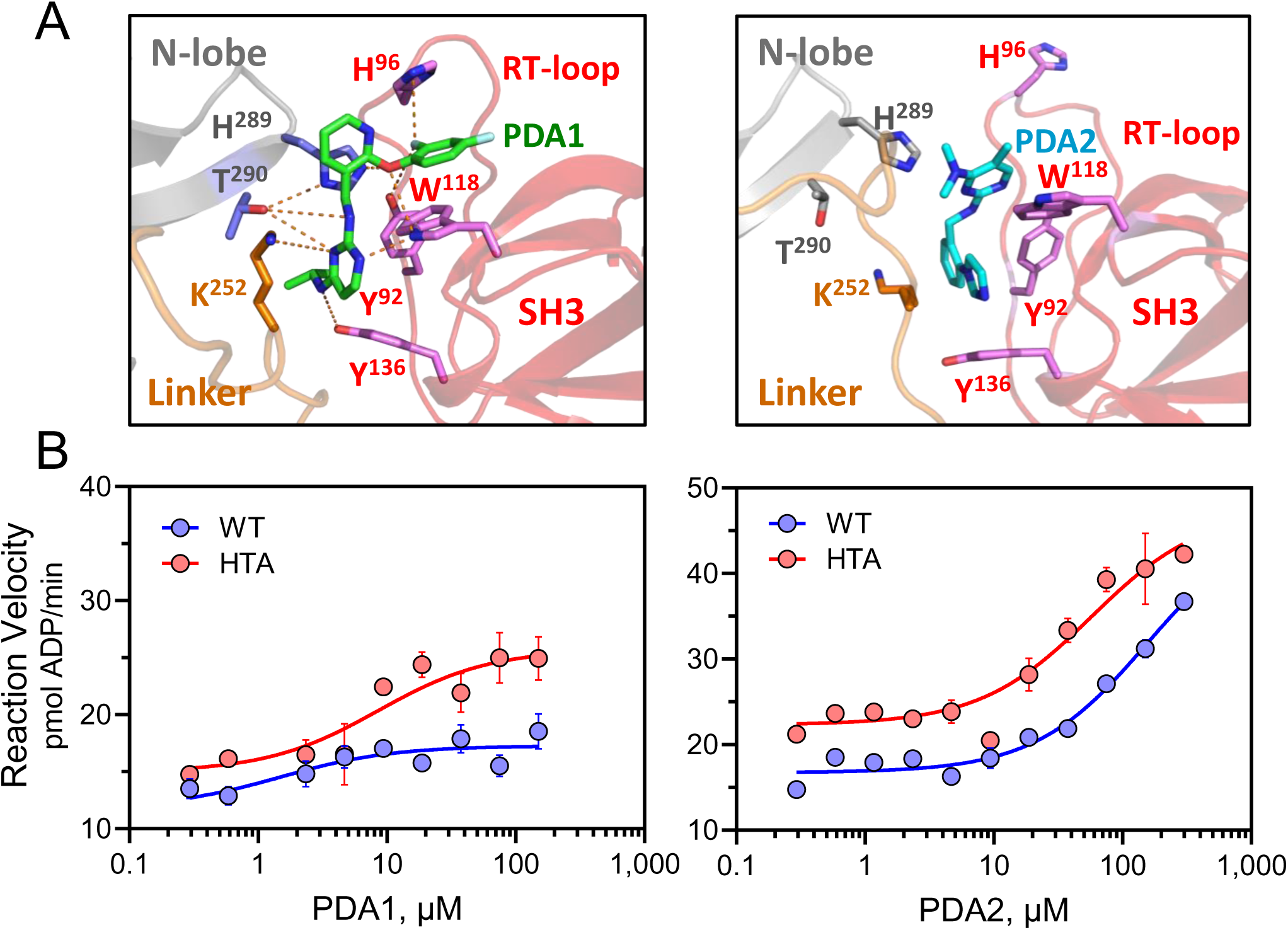
Molecular dynamics (MD) simulations of PDA ligand binding to near-full-length Hck and verification of the binding site by kinetic kinase assay. PDA1 and PDA2 were docked with the crystal structure of near-full-length Hck followed by 600 ns MD simulations as described under Materials and Methods. Movies of the complete simulations are provided in the Supplemental Information. A) Molecular models from the final frame of the simulation show that PDA1 forms a bridge between the SH3 domain and the N-lobe of the kinase domain while PDA2 perturbs the structure of the SH3 domain RT loop to disrupt the SH3-linker interface. B) Kinetic kinase assays were performed with recombinant Hck proteins over the range of PDA1 and PDA2 concentrations shown. The assays compared PDA effects on wild-type (WT) Hck vs. a double mutant in which kinase domain N-lobe residues H289 and T290 were replaced with alanine (HTA mutant). This mutant is based on the MD simulations which predicted a key role for these N-lobe amino acids in PDA1-mediated bridging with the SH3 domain as shown in part A.

To determine whether the N-lobe contacts predicted by MD simulation are involved in PDA1 stabilization of Hck, we produced recombinant near-full-length Hck with an H289A/T290A double mutation (Hck-HTA mutant). The effects of the PDA compounds on Hck-HTA activity were then assayed using the kinetic kinase assay. The baseline activity of Hck-HTA was somewhat higher than wild type, which may reflect loss of stabilizing contacts between the N-lobe and the linker (e.g., hydrogen bonding of Thr290 with Lys252 in the linker), though this mutation does not induce full activation of the kinase. Both PDA1 and PDA2 activated Hck-HTA (Figure 7B), unlike wild-type Hck, which is only activated by PDA2. These data support the MD simulation prediction that PDA1 requires N-lobe residues His289 and Thr290 to interact with Hck without disrupting the downregulated conformation.

### PDA1 binds WT and ATP-site inhibitor resistant mutant Hck in cells

To determine whether the PDA compounds affect Hck signaling in a cellular environment, we used the human TF-1 cell line as a model system. TF-1 erythroleukemia cells do not express endogenous Hck and require GM-CSF or other myelopoietic cytokines to proliferate in culture (37). Introduction of active forms of Hck transforms the cells to a cytokine-independent phenotype, allowing assessment of Hck-dependent signaling (38).

Using recombinant retroviruses, TF-1 cells were initially transduced with wild-type Hck or a fusion protein in which the 70-amino-acid coiled-coil oligomerization domain of Bcr was fused to the Hck N-terminus (cc-Hck). This cc-Hck fusion protein causes the protein to oligomerize and become activated like the Bcr-Abl oncoprotein in CML resulting in GM-CSF-independent growth and survival. Consequently, the cells become susceptible to ATP-site Hck inhibitors such as A-419259. Cells expressing wild-type Hck, on the other hand, still require GM-CSF and remain insensitive to this inhibitor. To confirm the cc-Hck phenotype, we assessed the A-419259 sensitivity of TF-1/cc-Hck cells in concentration-response experiments (Figure S7A). A-419259 treatment of TF-1/cc-Hck cells yielded an IC_50_ value of 7.75 ± 1.03 nM, while the parent cell line as well as cells expressing wild-type Hck showed no loss of viability at the highest concentration tested (1,000 nM). These results are consistent with previous findings that coiled-coil fusion stimulates Hck kinase activity and signaling that is inhibited by A-419259 treatment (38).

We next evaluated the effect of PDA1 and PDA2 on the viability of all three TF-1 cell populations. Cells expressing cc-Hck were sensitive to PDA1 treatment, with a IC_50_ value of 4.2 ± 0.57 µM (Figure 8A). However, cells expressing wild-type Hck as well as the parent cell line were similarly sensitive, with overall IC_50_ values of 4.8 ± 0.42 and 5.2 ± 0.68 µM, respectively, suggestive of off-target effects. Closer inspection of the concentration-response curves showed that PDA1 caused greater growth suppression between 0.3 and 3 µM in the TF-1/cc-Hck cells compared to the other TF-1 cell populations (Figure 8A). This concentration range offered a window to explore possible Hck-directed effects of PDA1.

**Figure 8.**
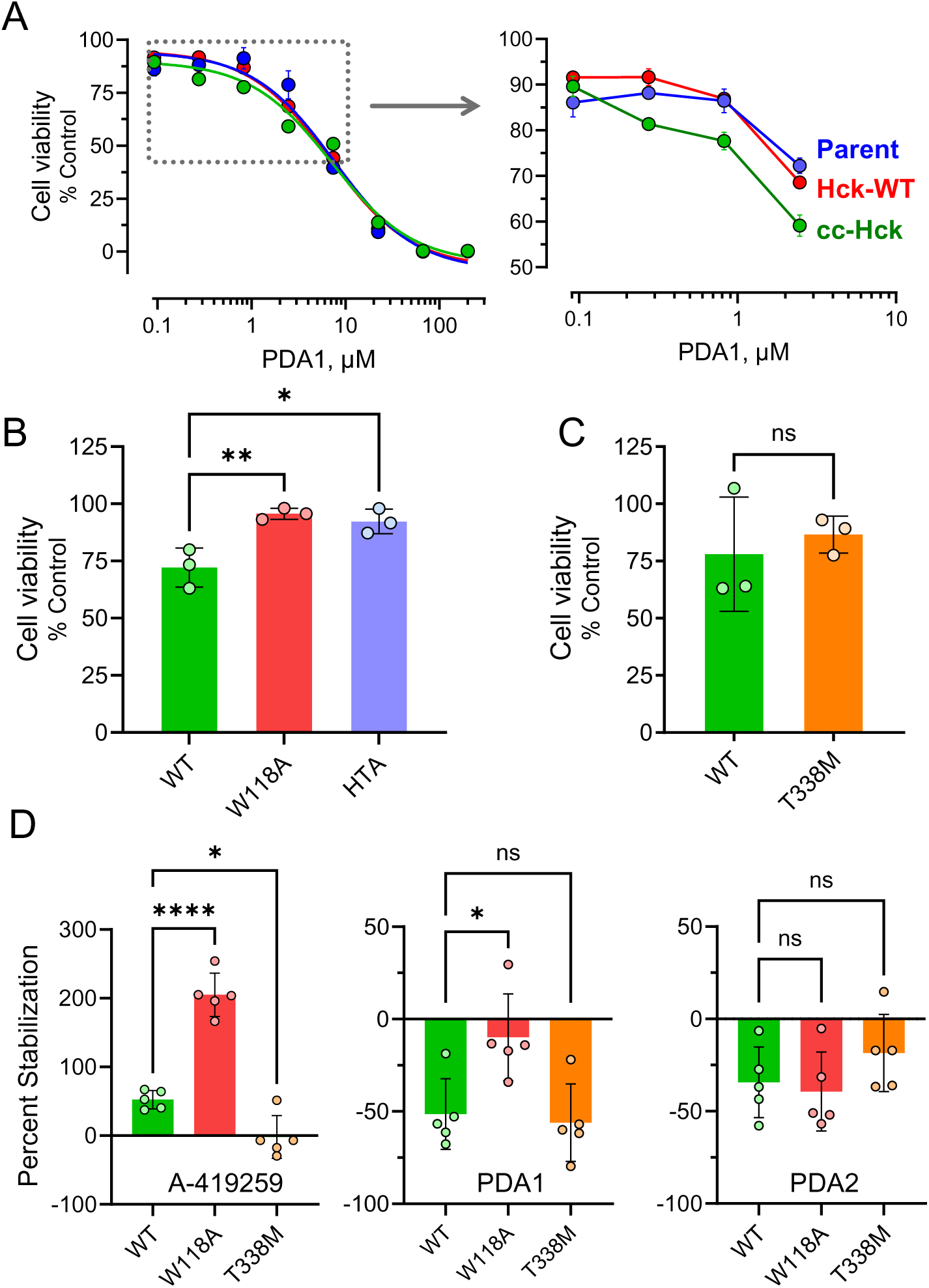
Interaction of PDA1 with Hck in TF-1 myeloid leukemia cells. A) TF-1 erythroleukemia cells were transduced with wild-type Hck (WT) or with an active coiled-coil fusion protein (cc-Hck). Cells were cultured in the presence of PDA1 over the range of concentrations shown or the DMSO carrier solvent alone (0.1%) as a control. After 96 h, cell viability was assessed using the CellTiter-Blue assay. Data were normalized to the DMSO control and each point represents the mean of three replicates ± SE. The curves were best-fit by non-linear regression and IC_50_ values are presented in the text. The boxed area of the main curve is expanded on the right to show enhanced sensitivity of cc-Hck cells to PDA1. B/C) TF-1/cc-Hck cells with wild-type (WT) kinase domains or mutations in the SH3 (W118A) or N-lobe (HTA), were treated with PDA1 (1 µM) and viability was assessed in triplicate assays 96 h later (B). These experiments were replicated to compare TF-1 cells expressing WT-cc-Hck protein and Hck with a mutation of the gatekeeper residue, T338M (C). Bar heights indicate mean cell viability relative to the DMSO control ± SE. Significant differences were determined by one-way ANOVA (B) or Student’s T-test (C). *, p < 0.05; **, p < 0.01; ns, not significant. D) Cellular thermal shift assay (CETSA). TF-1 cells were transduced with wild-type Hck or the SH3 (W118A) and gatekeeper (T338M) mutants. In-cell interaction of Hck with A-419259, PDA1, or PDA2 was assessed as the difference in Hck protein recovery by immunoblot following incubation at 35 °C vs. 48 °C and is plotted as the mean difference (stability) ± SE for five independent determinations. Significant differences were evaluated by Student’s t test. *, p < 0.05; ns, not significant.

To investigate this possibility, additional TF-1 cell populations were produced that express cc-Hck with the SH3-W118A mutation that does not bind PDA1 and as well as the HTA double mutation that no longer forms the N-lobe bridge with PDA1 as predicted by MD simulations. Each of these cell populations, along with TF-1/cc-Hck cells with the wild-type kinase domain, were treated with PDA1 at a concentration of 1 µM and viability was determined 96 h later. Cell populations expressing each mutant form of cc-Hck showed significant rescue of PDA1-induced cytotoxicity compared to wild-type cc-Hck cells (Figure 8B). In a final experiment, TF-1 cells were transformed with a gatekeeper mutant of Hck in which Thr338 in the kinase domain is replaced with methionine (T338M). This mutation activates the kinase and confers resistance to ATP-site inhibitors, including A-419259, and activates the kinase sufficiently to enable GM-CSF-independent growth. Cells expressing Hck-T338M were as sensitive to PDA1 as the cc-Hck cells with a wild-type kinase domain, suggesting that PDA1 could be used to target cells which are resistant to ATP-site inhibitors (Figure 8C). Unlike PDA1, PDA2 showed very low potency against all TF-1 cell populations, with IC_50_ values > 190 µM (Figure S7B).

To confirm that PDA1 interacts directly with Hck in TF-1 cells, we used a cellular thermal shift assay (CETSA). In this assay, interaction of a ligand with a target protein often affects its thermal stability making it either more or less aggregation prone at higher temperatures. For this experiment, we used populations of TF-1 cells expressing either wild-type Hck, Hck-W118A, or Hck-T338M without the coiled-coil fusion. Each cell population was incubated with the PDA analogs, A-419259 as a positive control, or 0.1% DMSO (vehicle control) for 1 hour. Cell aliquots were then incubated at either 35 °C or 47.5 °C for 2 minutes and the amount of protein in the soluble fraction was quantified by immunoblotting for Hck with Actin as loading control. The results are presented as the difference in the percent stability of Hck in the presence of each compound compared to the DMSO control (Figure 8D).

A-419259 treatment increased the thermal stability of both wild-type Hck and the SH3-W118A mutant but had no effect on Hck-T338M which does not bind A-419259. In contrast to A-419259, PDA1 treatment decreased the stability of both wild-type Hck and the T338M mutant. However, this effect was lost with the W118A mutant, which does not bind PDA1 *in vitro*. These results provide direct evidence that PDA1 enters cells and interacts with Hck in an SH3 domain-dependent manner and are consistent with the role of Trp118 in binding as observed by NMR. Finally, PDA2 treatment had a small negative effect on Hck stability in all three cell lines. This result suggests that PDA2 may show low cell penetration, which is consistent with its relatively low cytotoxicity compared to PDA1 (Figure S7). Alternatively, PDA2 may not have a significant impact on the thermal stability of Hck, precluding a readout in this assay.

## Discussion

In a previous study, we identified a series of pyrimidine diamines that bind to the regulatory domains of Hck (27), a Src-family kinase expressed in myeloid hematopoietic cells and potential target for AML therapy (18, 20, 39). Here we demonstrate that two of these compounds, PDA1 and PDA2, share a binding site on the Hck SH3 domain. Surprisingly, these analogs have opposing actions on the overall conformation of Hck in solution despite the shared SH3 domain binding site. MD simulations combined with site-directed mutagenesis strongly suggest that this difference is due to alternative binding modes, with PDA1 stabilizing and PDA2 perturbing the SH3-linker-N-lobe interactions required for kinase regulation.

We identified the binding site of PDA1 and PDA2 on the Hck SH3 domain by ^1^H-^15^N HSQC NMR and HDX-MS. Both compounds occupy the PPII helix-interacting site on the SH3 domain centered around Trp118. SPR experiments reported during the initial discovery of these compounds suggested that they could not bind the SH3 domain alone (27). However, the NMR results presented here indicate that the SH3 domain is sufficient for ligand binding. The lack of binding previously reported may have been the result of low SH3 domain immobilization, suboptimal orientation following covalent attachment of SH3 to the SPR biosensor, or the flow-based nature of SPR. Despite a shared SH3 domain binding site, PDA1 and PDA2 have opposing effects on overall Hck conformational dynamics by HDX-MS. PDA1 induced overall stabilization of Hck proteins consisting of the unique, SH3, SH2, and linker (U32L) as well as near-full-length Hck, as reflected in reduced deuterium uptake by most peptides derived from these proteins. Interestingly, binding of PDA1 to SH3 reduced deuterium update in the unique domain of the U32L protein, suggesting that these domains interact as shown by previous NMR studies (9). In contrast, PDA2 disrupted the overall kinase conformation, increasing deuterium uptake in most peptides including several at a distance from the SH3 binding site. These results are consistent with kinase activity assays, where PDA2 induced kinase activation while PDA1 was without effect.

Both PDA1 and PDA2 protected SH3 domain peptides in the truncated U32L protein from deuterium exchange, consistent with SH3 as the binding site for both compounds (Figure 5A). However, SH3 peptides from near-full-length Hck did not show protection from exchange with either PDA1 or PDA2 (e.g. SH3 peptide 109-135; Figure 6D). In the apo state of the near-full-length protein, the SH3 domain is bound to the SH2-kinase linker and therefore protected from exchange. Binding of either ligand to the SH3 domain displaces the linker but maintains protection of SH3 from deuterium uptake due to the shared binding site. As a result, there is little net difference in exchange for the SH3 domain in the context of near-full-length Hck. A similar result was observed following HDX-MS analysis of Hck in complex with the HIV-1 Nef protein, which also binds to the SH3 domain to activate the kinase via linker displacement (40). In this case, the SH3 domain is either bound to the linker or to Nef and is protected from deuterium uptake in either case (41). Previous HDX-MS studies of an Hck SH3-SH2-linker protein have also shown that the SH3 domain does not engage the linker in the absence of the kinase domain (2). Thus, PDA ligand binding results in protection of the SH3 domain from deuterium uptake in the U32L protein because it is not bound to the linker (Figure 5A).

Previous work has shown autophosphorylated Hck can be stimulated further by SH3 domain displacement under identical assay conditions (33), indicating that autophosphorylation alone does not cause full kinase activation. This effect was shown again in the present study with the activation of Hck by PDA2. Unlike PDA2, PDA1 did not stimulate kinase activity in vitro, which was surprising given previous reports that proteins or peptides that bind to the Hck SH3 domain result in kinase activation as observed with PDA2 (33, 35). MD simulations and HDX-MS suggest that PDA1 stabilizes the closed state of Hck by bridging residues in the SH3 domain with kinase domain N-lobe residues His289 and Thr290. Mutagenesis of these N-lobe residues to alanines increased basal kinase activity and rendered Hck susceptible to activation by PDA1. While PDA1 binding appears to maintain the closed conformation of the overall kinase based on the HDX data, it did not suppress kinase activity. However, future analogs of PDA1 with higher affinity may show direct inhibitory potential. Regardless, selective SH3-binders such as PDA1 may represent useful drug leads by preventing SH3-dependent substrate recruitment to disrupt oncogenic signaling. Consistent with this idea, an Hck SH3 domain mutant showed reduced oncogenic activity in rodent fibroblasts due in part to reduced recruitment and activation of the Stat3 transcription factor (29).

Cell-based studies revealed some promising insights into the potential of PDA1 as a scaffold for future SFK allosteric inhibitor development. Cellular thermal shift assays demonstrate that PDA1 enters cells and interacts directly with Hck in a manner dependent on the SH3 domain. Consistent with this observation, PDA1 treatment had a modest impact on the viability of leukemia cells that are dependent on active Hck for growth, and this effect also required an intact SH3 domain. While PDA1 did not consistently inhibit recombinant Hck activity in vitro, in cells it may target an active kinase conformation or interfere with SH3-dependent substrate recruitment as described above.

In summary, our study identifies PDA1 as a starting point for SFK allosteric inhibitor development for use in combination with an orthosteric inhibitor in a double-drugging approach to combat emergence of drug resistance in AML or other SFK-dependent cancers. This approach shows promise in CML where the combination of an allosteric inhibitor (asciminib) and an orthosteric inhibitor (nilotinib) of Bcr-Abl completely suppressed the emergence of drug resistance in an animal model of CML in vivo while either drug individually did not (25). The authors conclude that the complementary, non-overlapping resistance profiles of these two inhibitors were responsible for suppression of drug resistance. That said, some caution is needed, because CML cells can evolve that develop Bcr-Abl independence and no longer respond to tyrosine kinase inhibitor therapy (42). Cellular thermal shift assays showed that PDA1 binding to Hck was not affected by a kinase domain gatekeeper mutation that renders Hck resistant to the ATP-site inhibitor, A-419259. This observation suggests that PDA1 analogues may complement ATP-site inhibitors similarly to asciminib. Future development of PDA1 will focus on enhancing binding potency, SH3 domain selectivity, and allosteric inhibition of kinase activity.

## Experimental procedures

### Expression and purification of recombinant Hck proteins

The nucleotide sequences for the SFKs Hck, Lyn and Src were PCR-amplified and subcloned into pET-28a (near-full-length kinases; SH3-SH2-kinase-tail), pET-30a (unique-SH3-SH2-linker; U32L) or pET-21a (SH3, SH3-SH2, and SH3-SH2-linker) expression vectors (Millipore Sigma) using either standard restriction digestion and ligation or the Gibson Assembly procedure (New England Biolabs). Mutations were added to proteins using either the QuickChange II site-directed-mutagenesis kit (Agilent) or Gibson Assembly. The plasmids were used to transform One Shot BL21 (DE3) STAR chemically competent *E. coli* (Thermo Fisher) using provider’s instructions. To facilitate expression of near-full-length protein by maintaining the closed conformation, the cells were co-transformed with a pET-DUET plasmid containing the PTP1B phosphatase catalytic domain (amino acids 1-283) and the full length Csk kinase. Bacterial cells were grown to an OD_600_ of 0.6-0.8 at 37 °C in 1 L of TB medium with 30 µg/mL kanamycin (SH3, SH3-SH2, U32L) or 30 µg/mL kanamycin and 50 µg/mL carbenicillin (near-full-length). The incubation temperature was decreased to 18 °C and protein expression was induced by addition of IPTG to a final concentration of 0.2 mM overnight. Following induction, the cells were pelleted via centrifugation and either immediately lysed or snap frozen in liquid nitrogen and stored at -80 °C until use.

Cells were suspended in Ni-IMAC binding buffer [25 mM Tris-HCl, pH 8.3, 500 mM NaCl, 20 mM imidazole, 2 mM β-mercaptoethanol (BME), and 10% glycerol] supplemented with cOmplete EDTA-free protease inhibitor cocktail (Millipore Sigma) prior to lysis via microfluidizer. The soluble protein fraction was isolated by ultracentrifugation at 100,000 g for 1 h at 4 °C. The clarified lysates were then loaded onto a 5 mL HisTrap HP column (Cytiva) and eluted with Ni-IMAC containing 500 mM imidazole. Fractions with near-full-length Hck (human p59 isoform amino acids 84-524 with an N-terminal 6x-His tag and modified C-terminal tail with sequence YEEIP) were pooled and dialyzed against affinity chromatography binding buffer (25 mM HEPES, pH 7.5, 1 mM EDTA, 2 mM BME and 10% glycerol) at 4 °C. Dialyzed protein was loaded onto a 5 mL HiTrap Blue HP affinity column (Cytiva) and eluted with the same buffer containing 3 M NaCl. The U32L protein (p59 Hck amino acids 22-261 plus an LEHHHHHH C-terminal tag) was dialyzed against 25 mM Tris-HCl, pH 8.3, 30 mM NaCl, 1 mM EDTA, 2 mM BME and 10% glycerol and purified using a 5 mL HiTrap Q HP anion exchange column (Cytiva) by eluting in buffer with 1 M NaCl. For the final polishing step, both proteins were diluted into size exclusion chromatography (SEC) buffer (20 mM Tris-HCl, pH 8.3, 150 mM NaCl, 10% glycerol, and 2 mM Tris(2-carboxyethyl)phosphine) and run over a HiLoad 16/600 Superdex 75 pg preparative SEC column (Cytiva). None of the mutations presented in this paper affected the protein purification process. The purity and integrity of each protein was checked after each step using SDS-PAGE and Coomassie blue protein staining.

^15^N-labeled proteins for NMR include the SH3 domain (p59 Hck amino acids 80-144), SH3-SH2 (amino acids 75-246), and the SH3-SH2-linker (amino acids 84-261). All three domain proteins had an additional 6x-His C-terminal tail connected with a two-residue Leu-Glu linker. BL21 (DE3) STAR cells were transformed with the corresponding pET-21a expression plasmids and grown in M9 defined medium containing ^15^NH_4_Cl as a sole nitrogen source supplemented with 50 µg/mL carbenicillin. Expression was induced by the addition of 0.2 mM IPTG overnight at 18°C. Cells were lysed followed by the initial Ni-IMAC purification step as described above. Fractions containing the recombinant proteins were pooled and dialyzed against 25 mM sodium phosphate, pH 7.5, 10 mM NaCl, 0.02% sodium azide, and 2 mM BME. Proteins were loaded onto a 5 mL HiTrap Q HP anion exchange column and eluted using a gradient with the same buffer containing 1 M NaCl. The proteins were then dialyzed against NMR buffer (10 mM HEPES, pH 8.0, 100 mM NaCl, and 10 mM BME) followed by SEC on a HiLoad 16/600 Superdex 75 pg preparative column.

### ^1^H-^15^N HSQC NMR

All ^1^H-^15^N HSQC experiments with SH3, SH3-SH2 and SH3-SH2-linker proteins were performed at 30 °C in 10 mM HEPES buffer containing 100 mM NaCl and 10 mM 2-mercaptoethanol at pH 8.0 on Bruker 800 AVANCE spectrometer equipped with a 5 mm triple-resonance, z-axis gradient cryogenic probe. Final protein concentrations ranged from 60-70 µM as indicated in the figure legends. Data were processed and analyzed using NMRPipe (43) and CcpNmr Analysis (44), and plotted using nmrView (45). Backbone amide N-H signals were identified using previously published assignments for the Hck SH3-SH2-linker protein (28). Tryptophan indole resonances were assigned by comparing NMR spectra of Hck SH3-SH2 proteins with and without W118A or W119A mutations, and that of Hck SH3-SH2-linker protein with and without W254A. PDA1 and PDA2 titration experiments were performed using 50 mM stock solutions in 100% DMSO and diluted to a final concentration of less than 2% in the NMR samples. Control experiments showed minimal influence of DMSO on the HSQC spectra (Figure S3B).

### Surface plasmon resonance (SPR)

SPR analysis was conducted using a Reichert 4SPR instrument (Reichert Technologies). Proteins were immobilized on carboxymethyl dextran hydrogel biosensor chips (Reichert) using standard amine coupling chemistry with 1-ethyl-3-(-3-dimethylaminopropyl) carbodiimide hydrochloride (EDC) and N-hydroxysuccinimide (NHS). Proper folding of the proteins was assessed by flowing the SH3-binding VSL12 peptide (VSLARRPLPPLP) over the proteins in SPR buffer (10 mM HEPES, pH 7.4, 150 mM NaCl, 0.05% Tween 20, and 3 mM EDTA) over a range of concentrations. The PDA compounds were dissolved in DMSO to make 100 mM stocks which were diluted in PBS to a final DMSO concentration of 1%. Compounds were flowed over the immobilized proteins over a range of concentrations at 50 µL/min with a 90 s association phase and 120 s dissociation phase. The chip surface was regenerated with 1 mM NaOH after each compound injection. Each concentration of each compound was measured in quadruplicate, and the resulting sensorgrams were corrected for buffer effects. Kinetic and binding constants were calculated by fitting the sensorgrams using 1:1 Langmuir binding model in the TraceDrawer program (Reichert).

### Hydrogen-deuterium exchange mass spectrometry (HDX-MS)

HDX-MS was performed on the Hck regulatory region protein (unique-SH3-SH2-linker) and on near-full-length Hck (SH3-SH2-linker-kinase-phosphotail) in the absence and presence of PDA1 and PDA2. Each protein was diluted to 20 μM in equilibration buffer (20 mM Tris-HCl, pH 8.3, 100 mM NaCl, 3 mM TCEP, H_2_O). Each diluted protein (1 μL) was then combined with 1 μL of equilibration buffer containing 6% DMSO (control) or 1 μL of PDA1 or PDA2 (600 μM) in equilibration buffer with 6% DMSO (final Hck protein:PDA molar ratio of 1:30 with a final DMSO concentration of 3%). Hck:PDA and control mixtures were equilibrated at 23 °C for 1 hour and then placed on ice until labeling was initiated. Samples were diluted 18-fold with labeling buffer (20 mM Tris-HCl, pD 8.3, 100 mM NaCl, 3 mM TCEP, 300 µM PDA1 or PDA2, 3% DMSO, 99.9% D_2_O) at 23 °C, and labeling was quenched at various time points (10 s to 4 h) by addition of equal volumes of ice-cold quenching buffer (150 mM K_2_HPO_4_, pH 2.4, H_2_O). PDA compounds were included during the equilibration and deuterium labeling steps to maintain the same level of binding site occupancy prior to quench. Based on the K_D_ values obtained for each ligand with Hck using SPR, occupancy is estimated as 88% for PDA1 and 73% for PDA2. Quenched samples were injected immediately into a Waters M-Class HDX system (held at 0 °C) for online pepsin digestion (performed at 12 °C) and reversed-phase separation (46). Deuterium incorporation was measured with a Waters SYNAPT G2si HDMS in ion mobility mode equipped with standard ESI source. Data were analyzed with Waters Protein Lynx Global Server 3.0 and DynamX 3.0. All HDX MS measurements were performed in duplicate and are presented as the average (see HDX Supplementary Data file for additional details). The precision of deuterium level determination is ± 0.25 Da and therefore differences larger than 0.50 Da were considered significant.

### In vitro kinase assays

The activity of near-full-length Hck (wild-type or HTA mutant) with or without PDA compounds was measured using the ADP Quest kinetic kinase assay (DiscoverX/Eurofins) which fluorometrically follows the production of ADP (47). Kinase reactions (20 µL) were run in black 384-well plates in kinase assay buffer (15 mM HEPES, pH 7.4, 20 mM NaCl, 1 mM EGTA, 0.02% Tween-20, 10 mM MgCl_2_, and 0.1 mg/mL bovine γ-globulins). Hck proteins were first titrated with an excess of ATP (500 µM) and substrate peptide (YIYGSFK; 500 µM) relative to the published K_m_ values (33) to identify a kinase concentration (150 ng/well) that would allow detection of either inhibition or activation by the PDA compounds. ATP and the substrate peptide concentrations were then titrated to determine their respective K_m_ values, with subsequent experiments set near the K_m_ for both (30 µM). PDA compounds were initially solubilized in 100% DMSO and diluted in assay buffer to a working concentration of 3 mM in 3% DMSO. The PDA compounds were then serially diluted and added to the Hck proteins in the presence of all other assay components except for ATP. The final maximum PDA concentration assayed was 300 µM and all assay wells contained 0.3% DMSO. The assay plate was incubated for 15 min at room temperature to allow for PDA binding followed by addition of ATP to start the kinase reactions. Hck activity was measured with and without substrate peptide to determine total phosphorylation and autophosphorylation. Fluorescence was measured every 5 min for 3 h using a SpectraMax i3x plate reader (Molecular Devices). The fluorescence values for each condition were averaged across four technical replicates, and the values for autophosphorylation were subtracted from the total phosphorylation values to obtain the peptide substrate phosphorylation rates. For each compound concentration, the linear section of the reaction progress curve was fit by regression analysis (GraphPad Prism, version 10). The slope of the line and the calculated SEM for each concentration was then graphed as a function of compound concentration followed by nonlinear regression analysis to calculate EC_50_ values where possible. Fluorescence values were converted to pmol ADP using a standard curve of known ADP concentrations.

### Alignment of Src-family kinase amino acid sequences

Amino acid sequences of the eight human SFKs were downloaded from Uniprot and aligned using Clustal Omega (https://www.ebi.ac.uk/jdispatcher/msa/clustalo; EMBL) to determine the sequence identity of each member to Hck. The sequences for the SH3, SH2, and kinase domains of the proteins were taken as defined in their Uniprot entries. The ATP-binding site residues were defined as any amino acid which had at least one atom within 7.5 Å of AMP-PNP in the crystal structures of Hck or Src (PDB: 1AD5 and 2SRC, respectively). These domains were similarly aligned to determine identity to Hck.

### Molecular Dynamics Simulations

The structures PDA1 and PDA2 were docked to a recent co-crystal of Hck bound to A-419259 in the active site, which adopts the overall closed conformation shown in Figure 1A. To enable access to the SH3 binding site, the SH2-kinase linker was removed (residues 244 to 260), then compounds were docked using the program Smina (36) with default parameters. A search area was defined by residues on the SH3 domain which demonstrated CSPs in the presence of each PDA by HSQC NMR (Figure 2) plus an outer shell of 5 Å. Both compounds interacted with SH3 W118 and were thus redocked to a smaller region centered on this residue using higher exhaustiveness and a higher number of modes. Poses were then manually inspected prior to simulation.

All simulations were run using the Amber ff14sb force field for the protein, the general AMBER force field (GAFF) for the compounds, and tip3p water model (48). Additionally, Hck was simulated in the autoinhibited state with Y527 phosphorylated, the parameters for the phosphotyrosine were obtained from Homeyer et al. (49) The Amber function tleap was used to create an octahedral water box with periodic boundaries and at least 15 Å between protein atoms. Enough sodium and chloride ions were added to the edge of the box to reach a concentration of 150 mM. The system was minimized, heated to 300 K, and equilibrated to reduce unfavorable interactions. During minimization and heating, restraints on the protein and compounds were progressively weakened to zero for equilibration and production runs. Triplicate production runs of at least 300 ns were run for both compounds. Since the compounds were predicted to bind at the SH3-linker interface, generating starting structures from the poses with all Hck atoms and no clashes involved modeling potential motion of the SH3 and SH2 domains relative to the rest of the protein. To this end, the SH2 and SH3 domain region of Hck was simulated alone according to the above procedure. Then, the compound pose was aligned to the simulated SH3 domain. Finally, this complex was inspected for clashes and the RMSD (without fitting) of the simulated SH3 domain was ensured to be no more than 1.5 Å from the initial crystal structure. Three starting structures were generated and simulated for each compound to improve confidence and ensure the results are not dependent on a specific starting structure.

### TF-1 cell culture and viability assays

TF-1 cells were sourced from the American Type Culture Collection (ATCC) and cultured in RPMI 1640 medium supplemented with 10% fetal bovine serum (FBS), 100 units/ml of penicillin, 100 μg/ml of streptomycin sulfate, and 0.25 μg/ml of amphotericin B (Antibiotic-Antimycotic; Gibco/ThermoFisher). Parental TF-1 cells and transduced cell populations expressing wild-type Hck also require human GM-CSF (1 ng/mL; ThermoFisher) to proliferate and survive in culture. 293T cells were obtained from the ATCC and cultured in DMEM supplemented with FBS and prophylactic antibiotics as described above.

Full-length cDNA clones of wild-type Hck and cc-Hck were subcloned into the retroviral expression vector pMSCV-puro (Takara Bio). Mutations were introduced into the coding sequences using Gibson Assembly (New England Biolabs). Retroviral stocks were produced in 293T cells co-transfected with each pMSCV construct and an amphotropic packaging vector. For retroviral transduction, TF-1 cells (10^6^) were resuspended in 5.0 mL of undiluted viral supernatant and centrifuged at 1,000 × g for 4 h at 18 °C. Viral supernatants were supplemented with Polybrene (4 μg/mL; Millipore Sigma) to enhance viral infection. Forty-eight hours later, transduced cells were selected 3 μg/mL puromycin and maintained in 1 μg/mL puromycin. TF-1 cell populations expressing cc-Hck proliferated in a GM-CSF-independent manner and were subsequently sub-cultured in the absence of GM-CSF.

To assess viability, cells were seeded at 10^5^/mL in the presence or absence of inhibitors with DMSO as carrier solvent (0.1% final concentration). Cell viability was assayed using the CellTiter-Blue reagent and the supplier’s instructions (Promega). Fluorescence intensity, which corresponds to changes in viable cell number, was measured using a SpectraMax i3X microplate reader (Molecular Devices). Fluorescence measurements were corrected for background observed with wells containing media only. Each experiment included three technical replicates per condition and concentration-response data were best-fit by non-linear regression analysis to determine IC_50_ values (GraphPad Prism).

### Cellular thermal shift assay

This experimental procedure was adapted from Jafari *et al*. (50) TF-1 cells expressing full length wild-type or mutant Hck were cultured as described above to a density of 10^6^ cells/mL. To establish a baseline thermal shift curve, cells (1.3 x 10^7^) were transferred to a 10 cm^2^ cell culture plate and in medium containing 0.1% DMSO. Following incubation at 37°C for 1 h, 8 x 10^6^ cells were removed and pelleted at 300 x g for 4 min. Cells were washed with PBS, pelleted, and resuspended in 800 µL of PBS with cOmplete protease inhibitor cocktail (Millipore Sigma) for a final concentration of 10^7^ cells/mL. Cell aliquots (50 µL) were pipetted into 12 x 0.5 mL PCR tubes and placed on a PCR gradient heating block set for a gradient between 34.9 and 55.7 °C for 3 min. Cells were then incubated at room temperature for 3 min before snap freezing in liquid nitrogen. The samples were then stored at -80 °C or thawed on a 25 °C heating block for 2 min, vortexed, and then frozen again. After a second thaw and vortex, the lysates were transferred to 1.5 mL microcentrifuge tubes and spun at top speed for 20 min at 4 °C. Clarified lysates (15 µL) were mixed with 15 µL of 2x SDS-PAGE sample buffer and heated at 95 °C for 5 min. Proteins were separated by SDS-PAGE, transferred to nitrocellulose membranes, and immunoblotted with rabbit Hck and mouse Actin antibodies. Proteins were visualized with secondary antibodies conjugated to IR dyes and imaged using a LICOR Odyssey clx instrument. Hck bands were quantified for each time point, and 3 replicate experiments were averaged to create a melting curve for further experiments.

To test PDA and inhibitor effects on Hck stability, TF-1 cells expressing wild-type or mutant Hck proteins were diluted to 10^6^ cells/mL and incubated with 300 nM A-419259, 100 µM PDA1 or PDA2, or 0.1% DMSO as vehicle control. Five replicates of 1.7 mL were prepared for each condition. Cells were incubated for 1 h at 37 °C and 1.4 mL from each sample was transferred to a microcentrifuge tube and pelleted by centrifugation at 300 x g for 4 min. Cells were washed in PBS, re-pelleted, and then suspended in 140 µL of PBS with cOmplete protease inhibitor cocktail (MilliporeSigma). Each sample was then split into two 50 µL aliquots in PCR tubes. One was heated at 35 °C and the other at 47.5 °C for 3 min; a control experiment showed that the higher temperature did not affect Actin stability. Cells were then lysed and clarified lysates were analyzed by immunoblot as described above. Hck band intensities were divided by the Actin band intensities to correct for loading. The Hck:Actin ratios at 47.5 °C were then divided by those at 35 °C to determine the percent aggregation at 47.5 °C. Each sample was immunoblotted twice, and the percent aggregation for each sample was averaged between the two technical replicates. The data was then normalized to the average stability of the DMSO control samples.

## Data availability

All data are described in the manuscript and in the Supporting Information file. HDX MS data have been deposited to the ProteomeXchange Consortium via the PRIDE (51) partner repository with dataset identifier PXD054878.

## Supporting information

This article contains supporting information.

## Supporting information

HDX Supplemental Data File

## Supporting Information

### Synthesis and analytical characterization of PDA1 and PDA2

#### Synthesis of pyrimidine diamine 1 (PDA1)

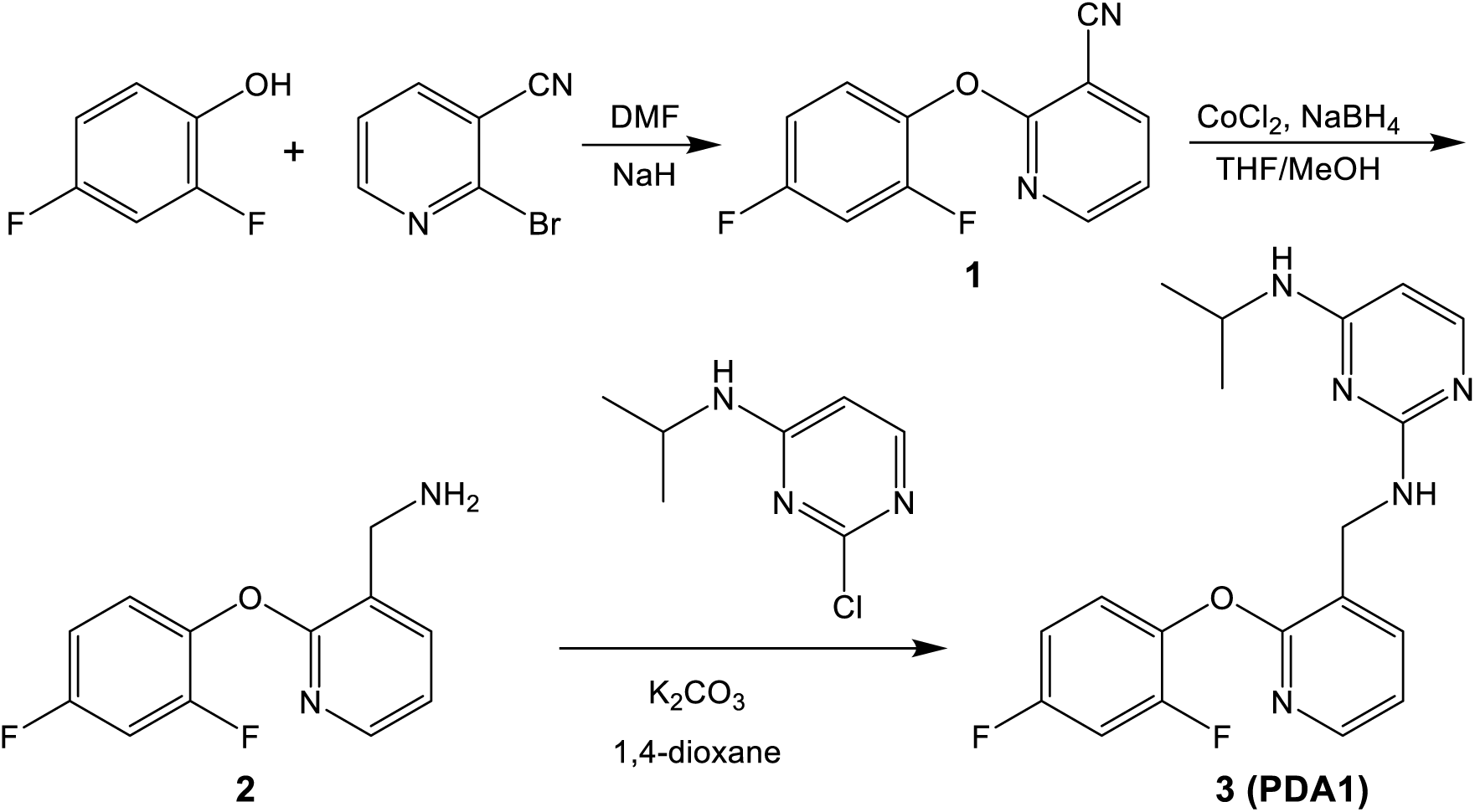

**2-(2,4-difluorophenoxy)nicotinonitrile, 1**. The phenol (0.746 g, 5.74 mmol) was dissolved in anhydrous DMF (30 mL) under Ar. NaH (60%, 0.262 g, 6.56 mmol) was added portion wise and and the mixture was stirred at room temperature for 1 h. The nitrile (1.000 g, 5.46 mmol) was added, the reaction stirred at room temperature for 2 h, and partitioned in EtOAc and water. The aqueous layer was extracted thoroughly with EtOAc. The organic layer was washed with water, brine, dried over Na_2_SO_4_ and solvents were evaporated to give a brown viscous residue. The residue was then purified by column chromatography on silica (ISCO-Rf, 0-50% EtOAc/hexanes) to give **1** as a white solid (0.719 g, 57%). ^1^H NMR (400 MHz, CDCl_3_): δ 6.93-7.00 (m, 2H), 7.14 (dd, 1H, J = 4.8 Hz, J = 7.2 Hz), 7.22-7.27 (m, 1H), 8.03 (dd, 1H, J = 2.0 Hz, J = 7.6 Hz), 8.29 (dd, 1H, J = 2.0 Hz, J = 5.2 Hz).

**(2-(2,4-difluorophenoxy)pyridin-3-yl)methanamine, 2.** A solution of nitrile **1** (0.250 g, 1.077 mmol) in THF/MeOH (1:1, 25 mL) was stirred at 0°C and CoCl_2_ (0.419 g, 3.230 mmol) portion wise followed by NaBH_4_ (0.407 g, 10.767 mmol). The mixture was stirred at 0°C for 20 min, filtered through Celite, and the Celite was washed thoroughly with dichloromethane. The organic layer was then treated with water and the water-organic mixture was filtered through Celite. The organic layer was separated, dried over Na_2_SO_4_ and the solvent evaporated to give a brown viscous residue. The residue was purified by silica chromatography (ISCO-Rf, 0-15% MeOH/dichloromethane) to yield **2** as a brown gel (0.150 g, 59%). LC-MS (ESI) m/z: calculated 236.22; observed 237.2 (M+1).

***N*2-((2-(2,4-difluorophenoxy)pyridin-3-yl)methyl)-*N*4-isopropylpyrimidine-2,4-diamine, 3 (PDA1).** The amine **2** (0.150 g, 0.635 mmol), anhydrous K_2_CO_3_ (0.263 g, 1.905 mmol), and 2-chloro-N-isopropyl-4-pyrimidinamine (0.163 g, 0.952 mmol) were combined with 1,4-dioxane (5 mL) in a sealed tube and heated to 135 °C for 16. The dioxane was evaporated and the residue purified twice by silica chromatography (ISCO-Rf, 0-15% MeOH/dichloromethane) to give **3** as a tan solid (0.051 g, 28%). ^1^H NMR (600 MHz, DMSO): δ 1.04 (s, 6H), 3.97 (s broad, 1H), 4.54 (d, 2H, J = 6.0 Hz), 5.70 (s, 1H), 6.75 (s broad, 1H), 6.95 (s broad, 1H), 7.07-7.15 (m, 2H), 7.37-7.44 (m, 2H), 7.62-7.66 (m, 2H), 7.90 (d, 1H, J = 3.0 Hz). ^13^C NMR (150 MHz, DMSO): δ 22.34, 104.91, 105.06, 105.09, 105.24, 111.52, 111.54, 111.67, 111.69, 119.10, 123.34, 125.20, 125.26, 137.10, 137.19, 144.28, 153.36, 153.45, 155.01, 157.96, 158.03, 159.41, 159.57, 159.65, 161.86, 162.03.^19^F NMR (500 MHz, DMSO): δ -123.73, -114.35.

#### Synthesis of pyrimidine diamine 2 (PDA2)

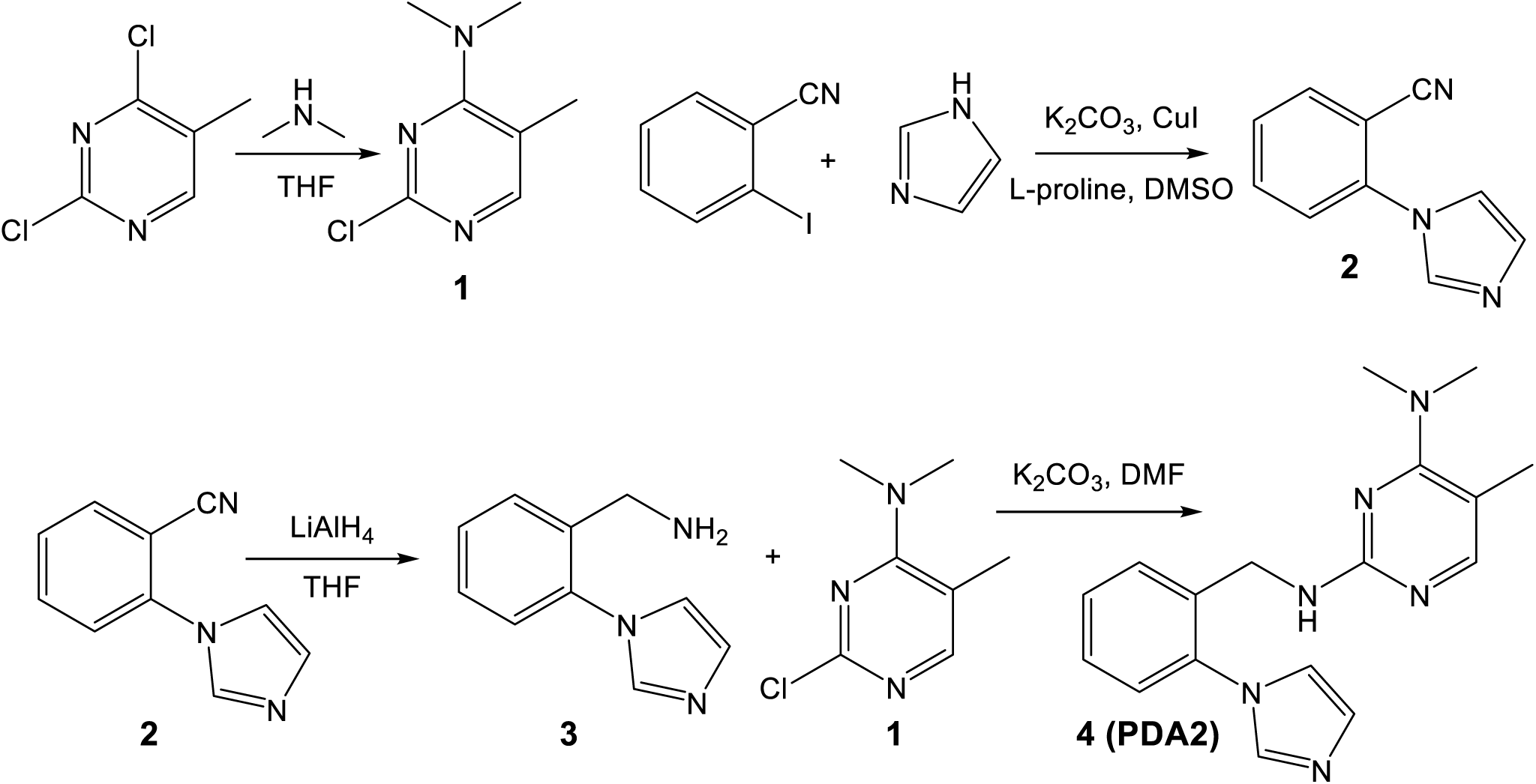

**2-chloro-N,N,5-trimethylpyrimidin-4-amine, 1.** 2,4-dichloro-5-methylpyrimidine (1.000 g, 6.135 mmol) was dissolved in THF (16 mL) under Ar. The reaction vessel was then cooled to 10 °C and dimethylamine (2M in MeOH, 6.6 mL, 13.3 mmol) was added and reaction was left to stir for 2.5 hours. Solvents were evaporated to give a white solid which was purified by silica chromatography (ISCO-Rf, 0-50% EtOAc/hexanes); 0.893 g, 85%. ^1^H NMR (600 MHz, CDCl_3_): δ 2.27 (s, 3H), 3.14 (s, 6H), 7.83 (s, 1H).

**2-(1H-imidazol-1-yl)benzonitrile, 2.** 2-iodobenzonitrile (1.000 g, 4.366 mmol), imidazole (0.357 g, 5.240 mmol), K_2_CO_3_ (1.208 g, 8.733 mmol), CuI (0.083 g, 0.437 mmol), and L-proline (0.101 g, 0.873 mmol) were combined in DMSO (10 mL) in a sealed vial and left to stir at 120 °C for 23 hours. The reaction was then partitioned in EtOAc and water and NH_4_OH was added to break the emulsion. The aqueous layer was extracted with EtOAc. The organic layer was washed with water and brine, and dried over Na_2_SO_4_ and solvents were evaporated to give a light yellow solid. The solid was then purified by silica chromatography (ISCO-Rf, 0-100% EtOAc/dichloromethane) to give the product as a white solid (0.358 g, 48%). ^1^H NMR (600 MHz, CDCl_3_): δ 7.28 (s, 1H), 7.37 (s, 1H), 7.47 (d, 1H, J = 7.8 Hz), 7.54 (t, 1H, J = 7.8 Hz), 7.75 (t, 1H, J = 7.8 Hz), 7.84 (d, 1H, J = 7.8 Hz), 7.87 (s, 1H).

**(2-(1H-imidazol-1-yl)phenyl)methanamine, 3.** A solution of nitrile **2** (0.150 g, 0.887 mmol) in THF (6 mL) under Ar was combined with LiAlH_4_ (2M in THF, 0.887 mL, 1.773 mmol) dropwise at 0 °C and the reaction mixture was stirred and allowed to come to room temperature over 1.5 h. The reaction was re-cooled back to 0 °C and quenched with NH_4_Cl dropwise until effervescence stopped. Na_2_SO_4_ was added and the reaction was left to stir at room temperature for 30 minutes. The reaction was then filtered through Celite, washed thoroughly with EtOAc/dichloromethane (1:1) and solvents evaporated to give an amber liquid. The liquid was purified by silica chromatography (ISCO-Rf, 0-15% MeOH/ dichloromethane) to give the product as a yellow residue (0.121 g, 79%). LC-MS (ESI) m/z: calculated (173.22); observed (M+1=174.2).

**N2-(2-(1H-imidazol-1-yl)benzyl)-N4,N4,5-trimethylpyrimidine-2,4-diamine, 4 (PDA2).** Amine **3** (0.121 g, 0.699 mmol), anhydrous K_2_CO_3_ (0.290 g, 2.096 mmol), and pyrimidine **1** (0.180 g, 1.048 mmol) were combined in DMF in a sealed vial and stirred at 140 °C for 15 h. DMF was evaporated, the reaction partitioned with water and MeOH/dichloromethane (1:10) and the aqueous layer extracted with EtOAc. The organic layer was washed with brine, dried over Na_2_SO_4_, and solvents evaporated to give a yellow liquid. The liquid was purified by silica chromatography (ISCO-Rf, 0-100% EtOAc/dichloromethane, 0-15% MeOH/dichloromethane) to yield 74 mg of a yellow viscous residue. The residue was repurified by silica chromatography (ISCO-Rf, 0-15% MeOH/dichloromethane containing 0.1% Et_3_N) to give a tan solid (0.018 g, 8%). ^1^H NMR (600 MHz, DMSO): δ 2.06 (s, 3H), 2.86 (s, 6H), 4.26 (d, 2H, J = 6.0 Hz), 6.92 (s broad, 1H), 7.10 (s, 1H), 7.29 (d, 1H, J = 7.8 Hz), 7.35 (t, 1H, J = 7.2 Hz), 7.41 (t, 1H, J = 7.2 Hz), 7.44 (s, 1H), 7.49 (d, 1H, J = 7.2 Hz), 7.57 (s, 1H), 7.85 (s, 1H). ^13^C NMR (150 MHz, DMSO): δ 7.14, 17.17, 51.94, 121.19, 126.08, 127.26, 128.18, 128.37, 128.72, 135.40, 136.45, 137.85, 158.12, 160.07, 164.11. LC-MS (ESI) m/z: calculated (308.39); observed (M+1=309.4).

**Figure S1:**
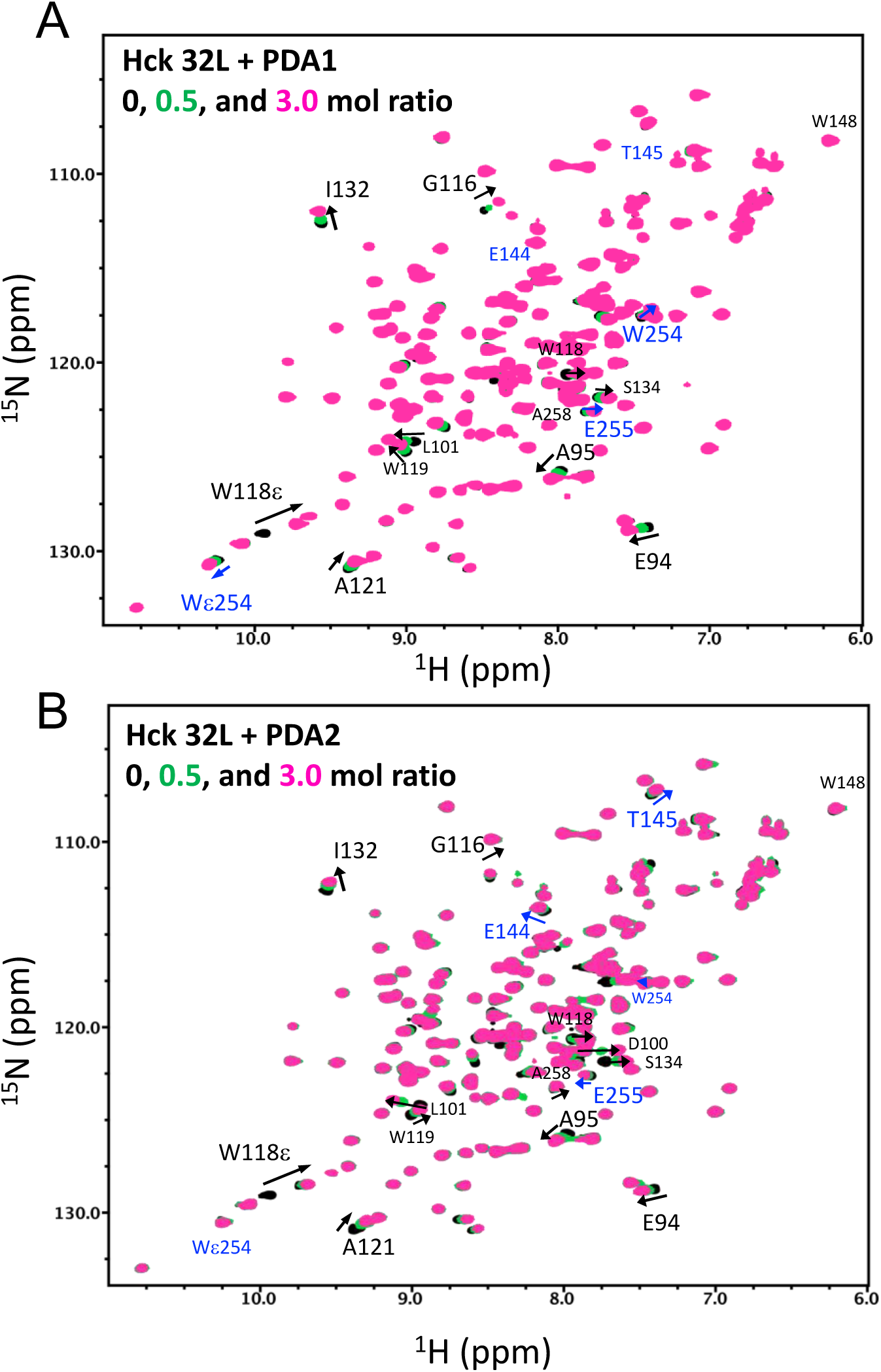
Complete HSQC NMR spectral changes of the Hck SH3-SH2-Linker protein upon PDA1 and PDA2 titration. An ^15^N-labeled Hck SH3-SH2-linker protein (60 µM) was titrated with (A) PDA1 or (B) PDA2, at protein:ligand ratios of 1:0 (black), 1:0.5 (green) and 1:3 (pink) and ^1^H-^15^N HSQC spectra were recorded. In both panels, chemical shift perturbations (CSPs) were observed in a subset of ^1^H-^15^N resonances (highlighted by arrows), indicating the specific interaction of the compounds with the protein. Most of the N-H amide resonances that exhibit significant CSPs localize to the SH3 domain (black arrows). PDA1 induced a CSP of the Trp254 indole resonance in the linker region but did not influence residues in the connecter region between SH3 and SH2. PDA2, on the other hand, induced CSPs of connector amides including Glu144 and Thr145, but not Trp254 in the Linker region (blue arrows). These results suggest that the shared pyrimidine diamine core in PDA1 and PDA2 interacts mainly with the SH3 domain while unique moieties in each compound affect the interdomain regions.

**Figure S2:**
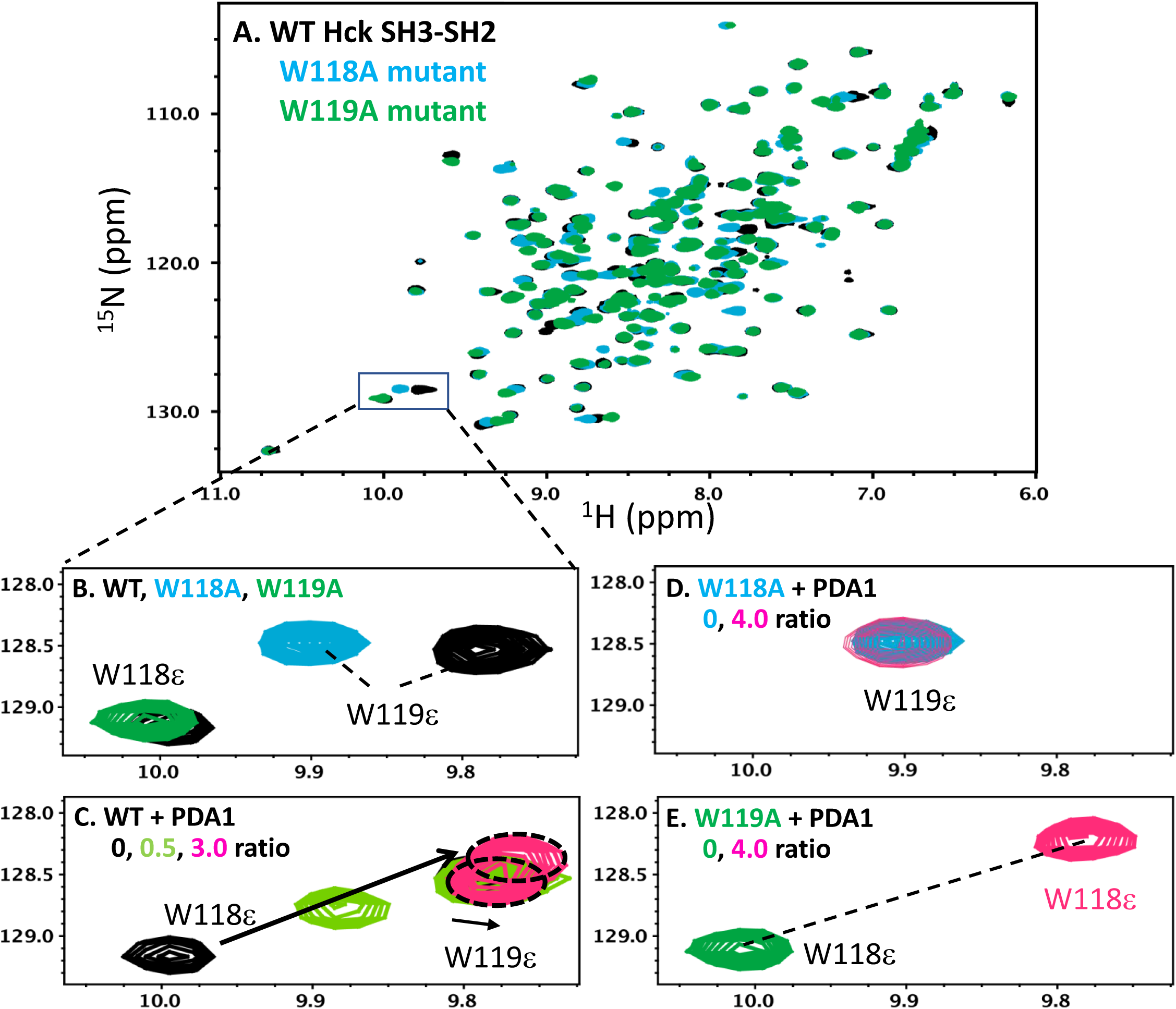
Assignment of indole N-H resonances of Hck SH3-SH2 Trp118 and Trp 119. A) Overlay of ^1^H-^15^N HSQC spectra of wild-type Hck SH3-SH2 (WT; black), the W118A mutant (cyan) and the W119A mutant (green), at Hck protein concentrations of 60 µM. B) Enlarged view of the tryptophan N-H indole (ε) region. C) Confirmation that the W118ε, and not the W119ε, resonance is significantly shifted upon PDA1 titration. D) Hck SH3-SH2 W118A mutant titration with PDA1, demonstrating no change of the shift of adjacent residue, W119. E) Titration of the SH3-SH2 W119A mutant with PDA1 exhibits a significant chemical shift in W118ε like wild type. Note that PDA2 also exhibited similar changes to W118ε (not shown). These observations assign the W118ε and W119ε chemical shifts in Hck SH3-SH2 and confirm that the SH3 W118 indole ring is critical for both PDA1 and PDA2 interaction. Note that the assignment of tryptophan indole resonances is consistent with that of a homologous Fyn SH3-SH2 protein.^1^

**Figure S3:**
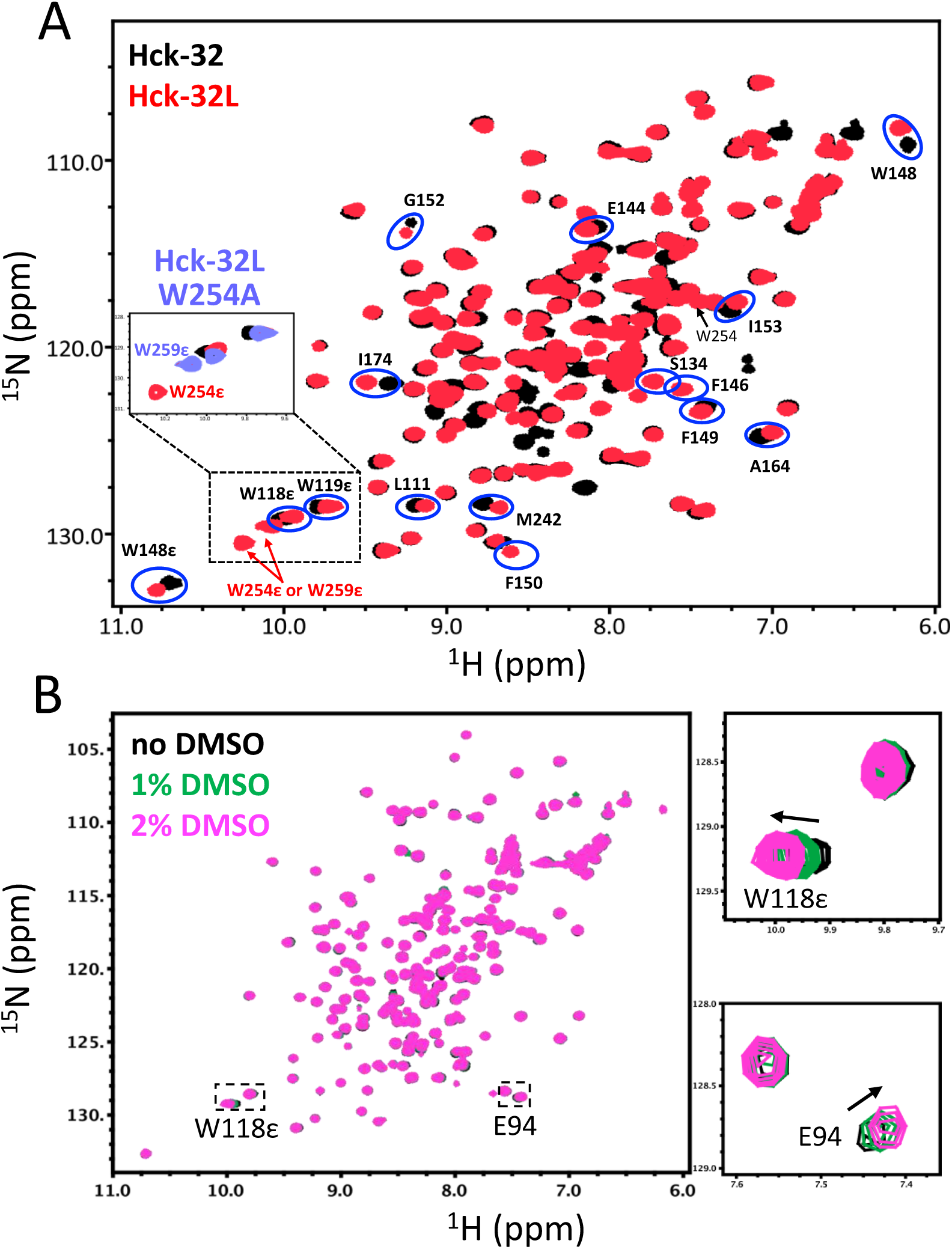
Linker effect on SH3-SH2 chemical shifts, assignment of linker W254 indole resonance, and DMSO control. A) Labeled residues exhibited significant chemical shift differences between the Hck SH3-SH2 (Hck-32; black) and SH3-SH2-linker (Hck-32L; red). The assignment of the backbone amides of Hck-32L is based on those of Jung *et al*.^2^ Two tryptophan residues are present in the linker region, W254 and W259. The tryptophan indole N-H resonances (ε) were assigned by comparing the NMR spectra of the wild-type and W254A mutant forms of the Hck-32L protein (**light blue** resonances in the box). Several residues in the connector between SH3 and SH2 exhibit different chemical shifts between Hck-32 and Hck-32L (circled in blue), suggesting that the linker interacts with the SH3-SH2 region in solution. B) DMSO effect on NMR data was assessed using Hck-32. The overall ^1^H-^15^N HSQC spectral features of Hck-32 were not altered even at 2% DMSO. A few minor chemical shift perturbations (CSPs) were observed for some residues (e.g., W118 indole N-H and backbone amide of Asp94, right panels.) However, the directions of these small CSPs were opposed to those of observed with the PDA1 and PDA2 titrations, and thus do not affect the PDA titration results. Protein concentrations of Hck-32, Hck-32L, and Hck-32L-W254 were 60 µM, 60 µM, and 70 µM, respectively.

**Figure S4:**
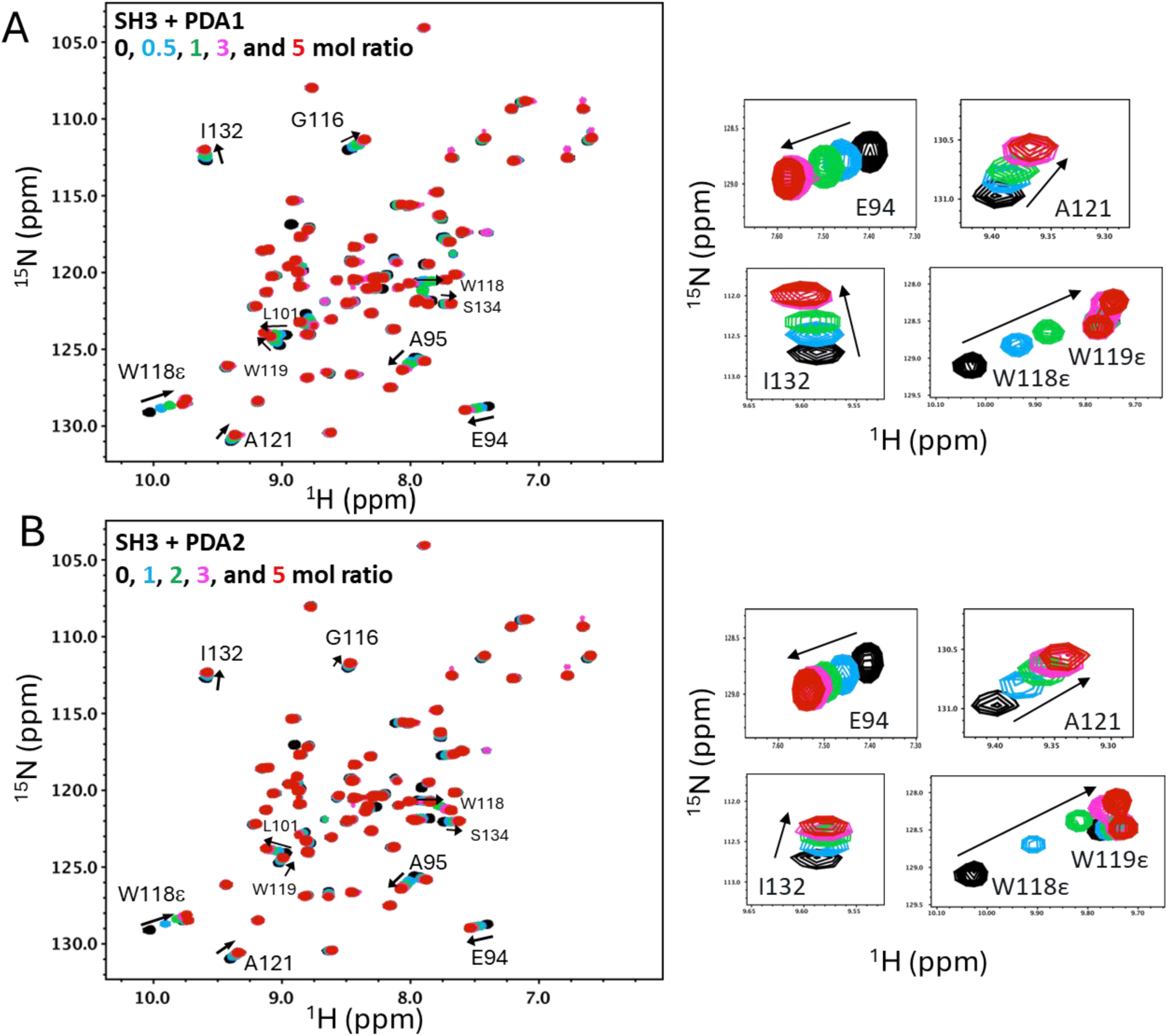
^1^H-^15^N HSQC NMR spectral changes of the Hck SH3 protein upon PDA1 and PDA2 titration. An ^15^N-labeled Hck SH3 protein (70 µM) was titrated with (A) PDA1 and (B) PDA2 at the molar ratios indicated. In both cases, chemical shift perturbations (CSPs) were observed in a subset of resonances (highlighted by black arrows), demonstrating a specific interaction of the compound with SH3 protein. The most substantial CSPs are enlarged on the right. These CSPs were essentially the same as those observed in the SH3 domain following PDA titration in the SH3-SH2-linker protein.

**Figure S5:**
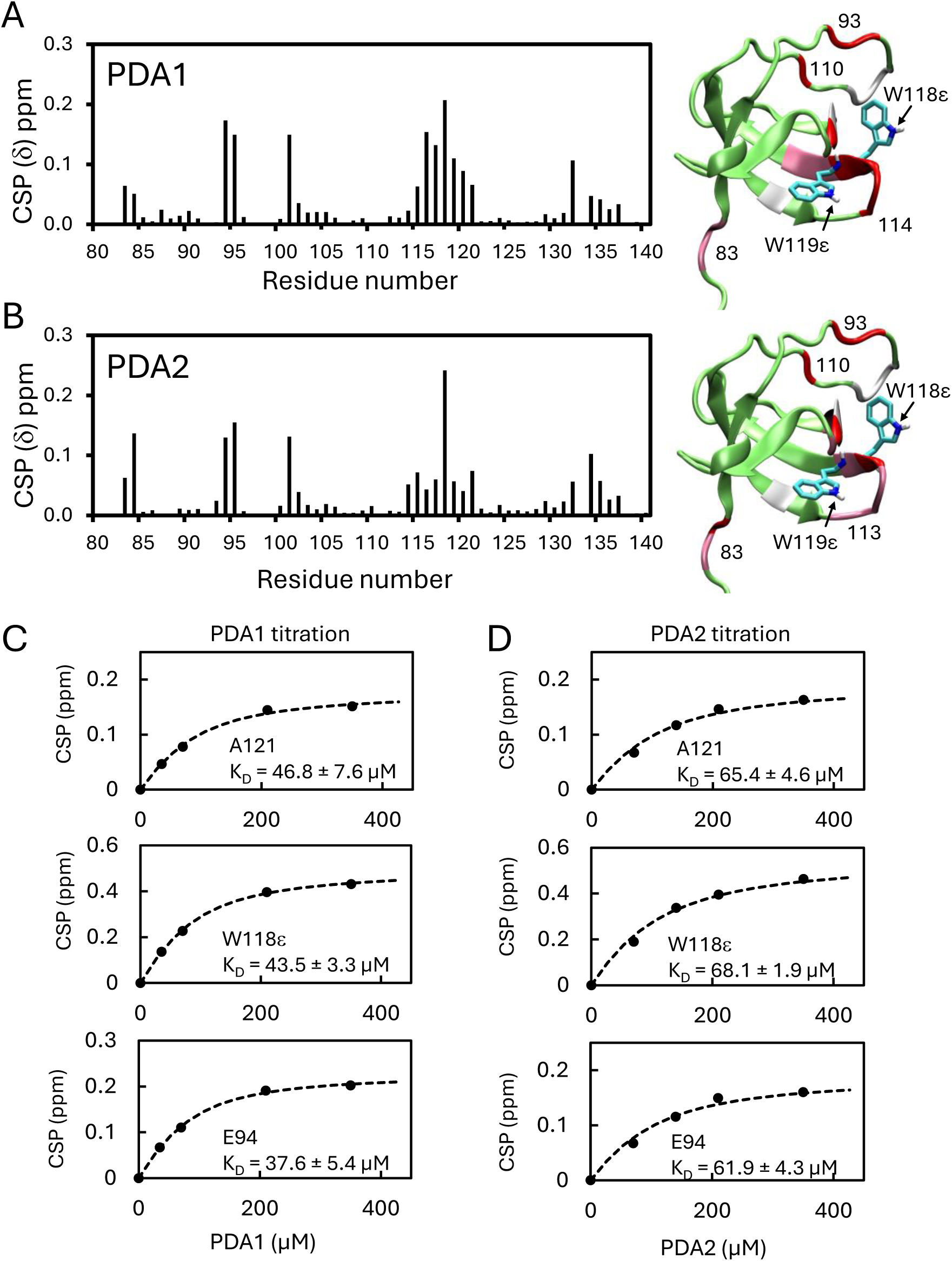
PDA1 and PDA2 NMR titration analysis with the Hck SH3 domain. Plots of CSPs at 1:3 SH3 protein:PDA ratio for A) PDA1 and B) PDA2 (SH3 concentration, 70 µM). CSPs are mapped on a ribbon structure of the Hck SH3 domain (*right*). Red, CSPs larger than one standard deviation above the average; pink, CSPs above the average; green, CSPs lower than the average; white, unassigned residues. CSPs were calculated using a combined shift of ^1^H and ^15^N with a weighting factor of 0.14 as per Williamson.^3^ Titration curves for three selected residues (Glu94 and Ala121 backbone amides; Trp118 indole amide) with C) PDA1 and D) PDA2. All three resonances show saturation with average dissociation constants of 42.6 ± 4.7 µM (PDA1) and 65.1 ± 3.1 µM (PDA2).

**Figure S6A:**
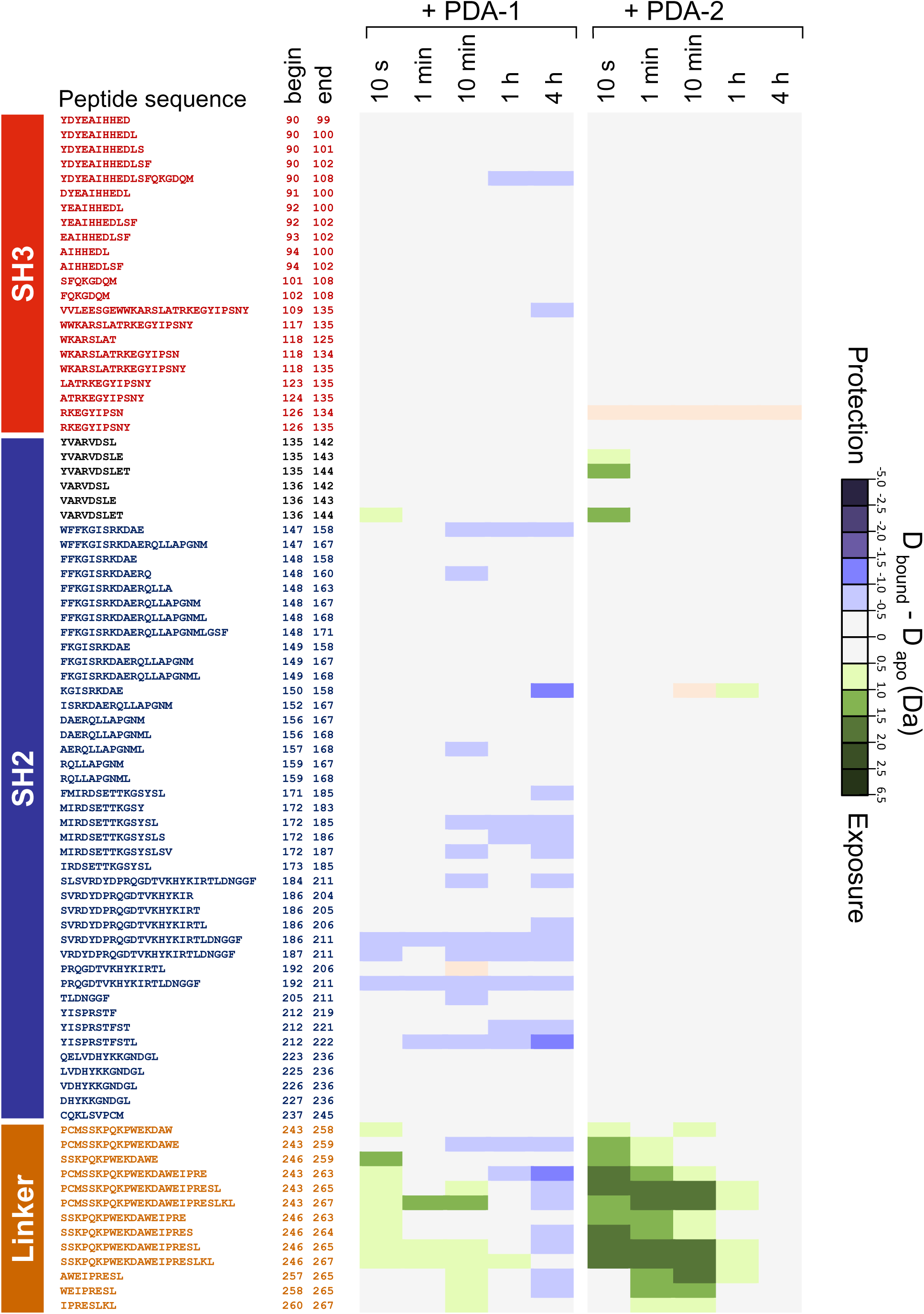
HDX-MS difference maps for near-full-length Hck in the presence of PDA1 and PDA2. Data are shown for the SH3 and SH2 domains plus the SH2-kinase linker. Kinase domain peptides are shown in Figure S6B.

**Figure S6B:**
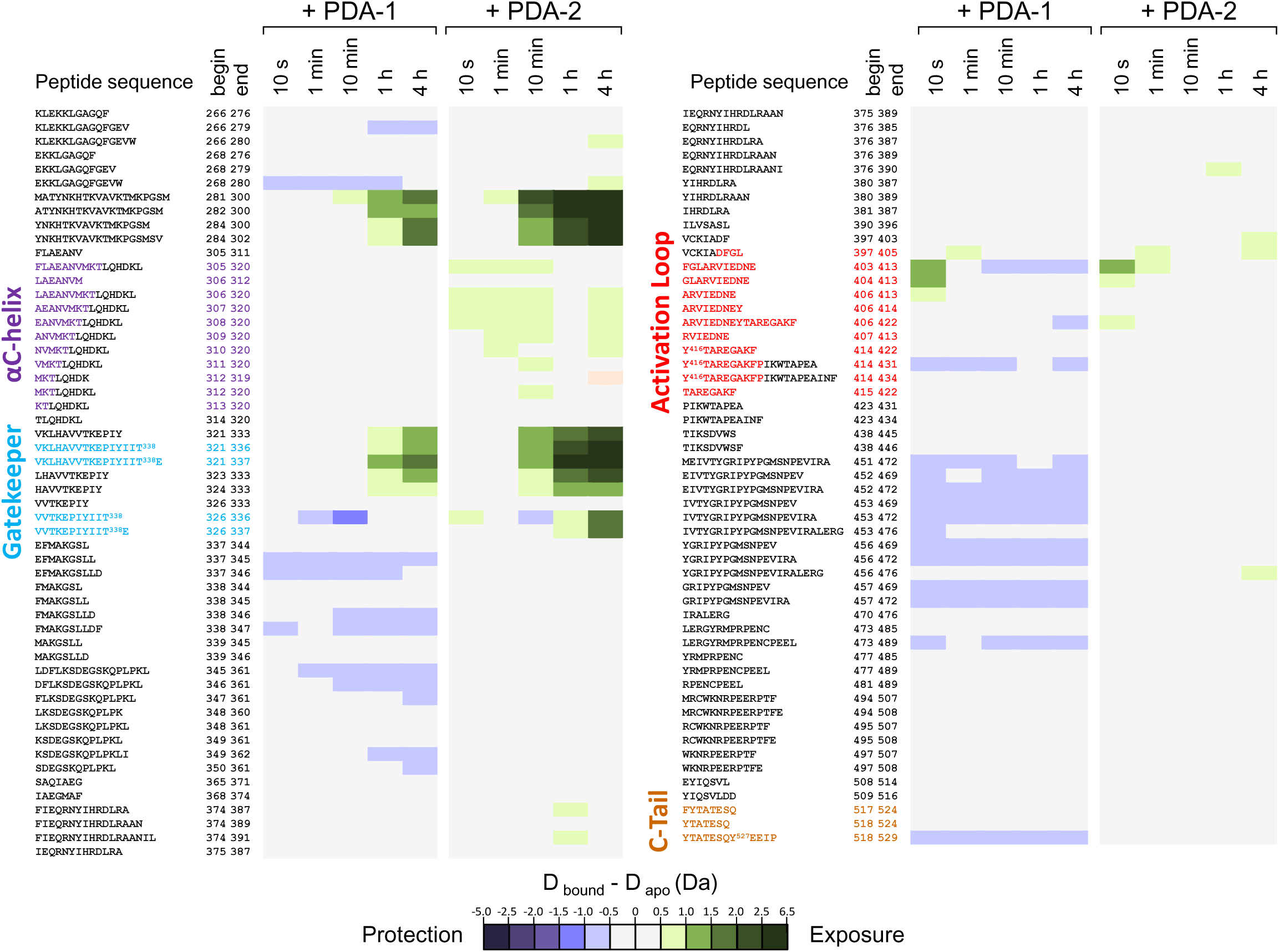
HDX-MS difference maps for near-full-length Hck in the presence of PDA1 and PDA2. Data are shown for the kinase domain; SH3 and SH2 domains plus the SH2-kinase linker peptides are shown in Figure S6A. Peptides containing residues from the **αC-helix**, the **gatekeeper residue (Thr338)**, the **activation loop and autophosphorylation site (pTyr416)**, and the **C-terminal tail and negative regulatory phosphotyrosine (pTyr527)** are highlighted.

**Figure S7:**
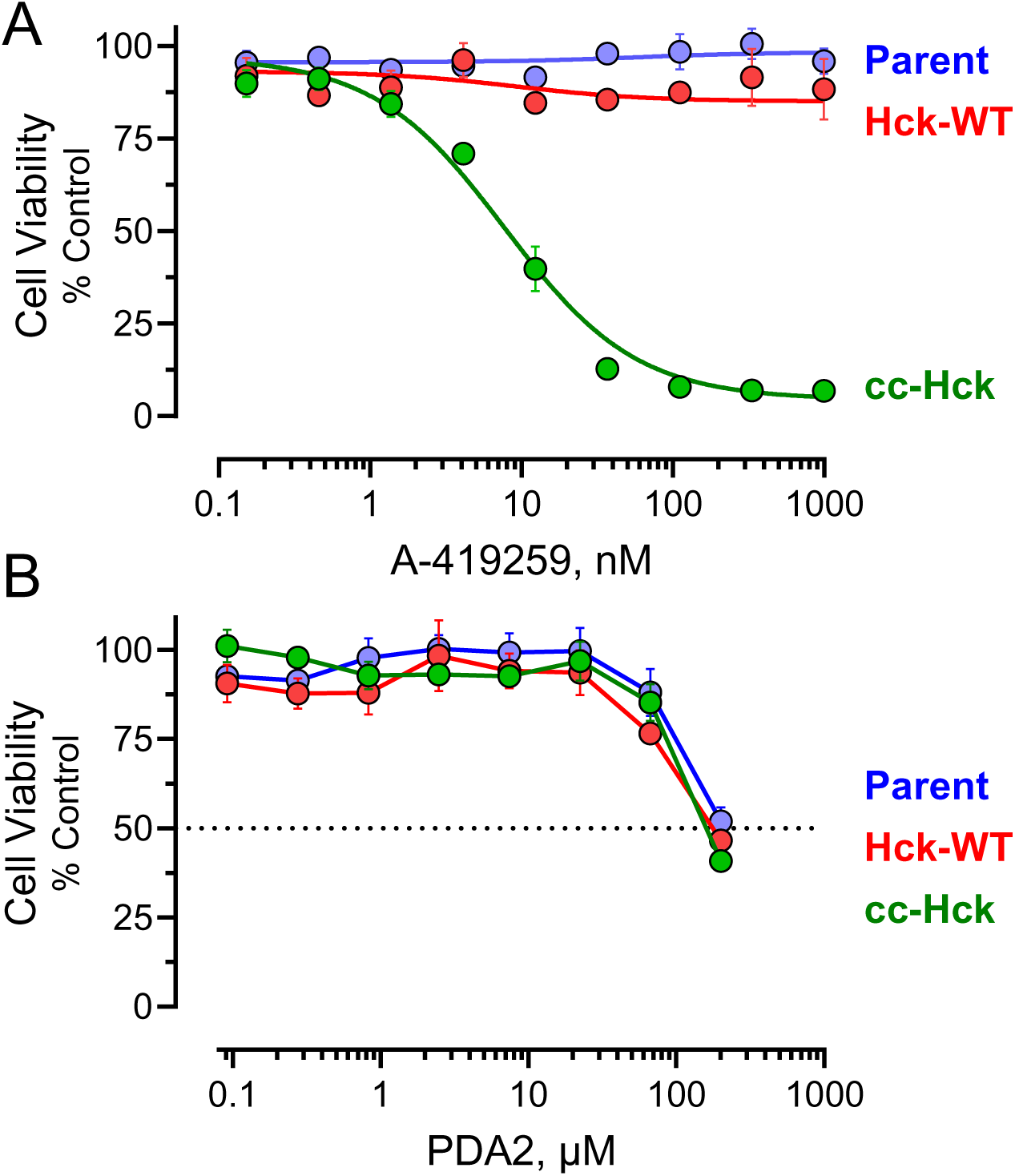
Effects of A-419259 and PDA2 on the viability of TF-1 cells expressing wild-type Hck and cc-Hck. Parental TF-1 cells or cell populations expressing wild-type Hck or an active cc-Hck fusion protein were incubated over a range of inhibitor concentrations or the DMSO carrier solvent (0.1%) alone as control. Parental cells and cells expressing wild-type Hck required GM-CSF for growth, while the TF-1/cc-Hck population acquired a cytokine-independent phenotype. Cell viability was determined 72 hours after addition of small molecules or vehicle using the CellTiter Blue cell viability assay (Promega). Results were normalized to the DMSO control values and are presented as mean percent control ± SE for triplicate determinations. A) Effects of orthosteric (ATP-site) inhibitor A-419259. B) Effects of PDA2, which is about 100-fold less potent than PDA1 (see main Figure 8) and shows no preference for any of the cell populations.

